# Pathogenic IgE-fated memory B cell responses retain functional plasticity

**DOI:** 10.1101/2023.11.28.567094

**Authors:** Kelly Bruton, Allyssa Phelps, Atai Ariaz, Allison Fang, Tina D. Walker, Jianping Wen, Sharon S. Khavkine-Binstock, Danielle Della Libera, Olivia Mann-Delany, Niels Peter H. Knudsen, Siyon Gadkar, Emily Grydziuszko, Joshua F. E. Koenig, Aidan Gagnon, Susan Waserman, Peter S. Andersen, Manel Jordana

## Abstract

Long-lived immunoglobulin (Ig) E responses against innocuous environmental and dietary antigens (Ags) are maintained by an IgG1-dominant memory B cell (MBC) compartment primed for IL-4 responsiveness. The plasticity of the MBC compartment destined for IgE class switch recombination (CSR), however, remains poorly understood. In this work, we report a critical IL-4/IL-13 dependency for the pathogenic IgE fate of type 2-polarized MBCs. Initiating a recall response in the absence of IL-4/IL-13 signaling diminished the type 2 MBC phenotype in mice and humans and, in mice, permitted the emergence of long-lived Ag-specific IgG2c^+^ MBCs. The divergence to a type 1-like response was dependent on IFN-γ signaling and arose from both unswitched and class-switched Ag-specific B cells *in vivo*. This reprogrammed fate was sustained even beyond therapeutic intervention, revealing fundamental insight into the plasticity of the allergen-specific MBC response.

**One Sentence Summary:** B cell responses to allergens can be reprogrammed away from a pathogenic fate through IL-4/IL-13 signaling blockade.

## INTRODUCTION

Classical immunological memory is marked by the propensity for rapid and optimized responses upon re-exposure to the target antigen (Ag). Within the B cell compartment, immunological memory is distinguished by high-affinity binding to Ag, a class-switched constant region conferring specialized effector functions, and altered cellular processes (*e.g.,* metabolism). Following vaccination, for example, viral exposure rapidly reactivates memory B cells (MBCs) facilitating plasma cell differentiation and boosting of neutralizing IgG titers, lessening the opportunity for infection (*1–3*). MBC reactivation, however, is pathogenic when perpetuating inflammatory responses to innocuous Ag, as occurs in allergic disease.

Allergies categorized as type I hypersensitivities are mediated by IgE. Humoral IgE responses are relatively short-lasting, yet reactivity to some allergens (*e.g.,* peanuts) can persist for a lifetime (*4–8*). Central to this persistence are IgG1^+^ MBCs which undergo sequential class-switching to replenish the transient IgE plasma cell pool (*5*, *9–12*). Likewise, it has been identified that IgE-fated MBCs, termed MBC2s, exhibit an IL-4-responsive phenotype marked by high expression of CD23 (FcεRII) and IL-4 receptor α (IL-4Rα) and low CD32 (FcγRII) expression (*13–15*). Indeed, seminal work on IgE established IL-4 as a critical molecule for IgE class switch recombination (CSR) upon primary sensitization (*16*, *17*). Despite the formation of a more specialized MBC compartment, IL-4 continues to be required for secondary IgE responses (*18–20*). Beyond inhibition of IgE CSR, loss of IL-4 signaling upregulates IFN-γ production suggestive of plasticity within the T cell compartment (*18*). Whether a parallel degree of plasticity exists in the Ag-specific MBC compartment, or if the commitment is irreversible, remains to be elucidated.

In this study, we identified a previously unrecognized plasticity of the MBC response within type 2 immunity. By inhibiting IL-4 signaling in a recall response, the pathogenic MBC2 phenotype was diminished, and upregulation of IFN-γ supported the emergence of long-lived IgG2c responses. Even beyond transient IL-4Rα blockade, IgG2c reemerged upon Ag exposure, suggesting some degree of stability in the type I-like reprogrammed antibody (Ab) response despite discontinued IL-4Rα blockade. Ag-specific IgG2b/c responses appeared to derive from both unswitched B cells and class-switched MBCs, where the addition of a type 1 adjuvant could augment the redirection of IgG1 memory to IgG2b/c. Collectively, our findings elucidate a malleable fate of the pathogenic IgE-fated MBC response and highlights the potential to therapeutically reprogram refractory lifelong allergies.

## RESULTS

### IL-4Rα signaling augments secondary germinal center activity

To assess the impact of *in vivo* IL-4Rα blockade (A4RA) on the B cell compartment in type 2 immunity, we employed a well characterized model of epicutaneous sensitization to ovalbumin (OVA). This strategy establishes a humoral OVA-IgG1 response and IgG1^+^ MBCs which, upon subsequent non-sensitizing subcutaneous (s.c.) re-exposures, facilitate IgE production and IgE-mediated clinical reactivity (*21*). With temporal control over the IgE response, we treated sensitized mice with A4RA or an isotype control (IC) prior to s.c. re-exposures, as previously described (*18*). Draining lymph nodes (dLNs) and spleens were collected for single-cell RNA-sequencing (scRNA-seq), single-cell VDJ-sequencing (scVDJ-seq), and flow cytometry, and blood was collected for immunoglobulin (Ig) analysis (**Fig. 1A**). For scRNA- and scVDJ-seq experiments an additional group of mice were administered cholera toxin (CT) and OVA (s.c.) to model a classical type 2 immunization strategy that yields robust humoral IgE responses and IgG1^+^ MBCs.

**Fig. 1.**
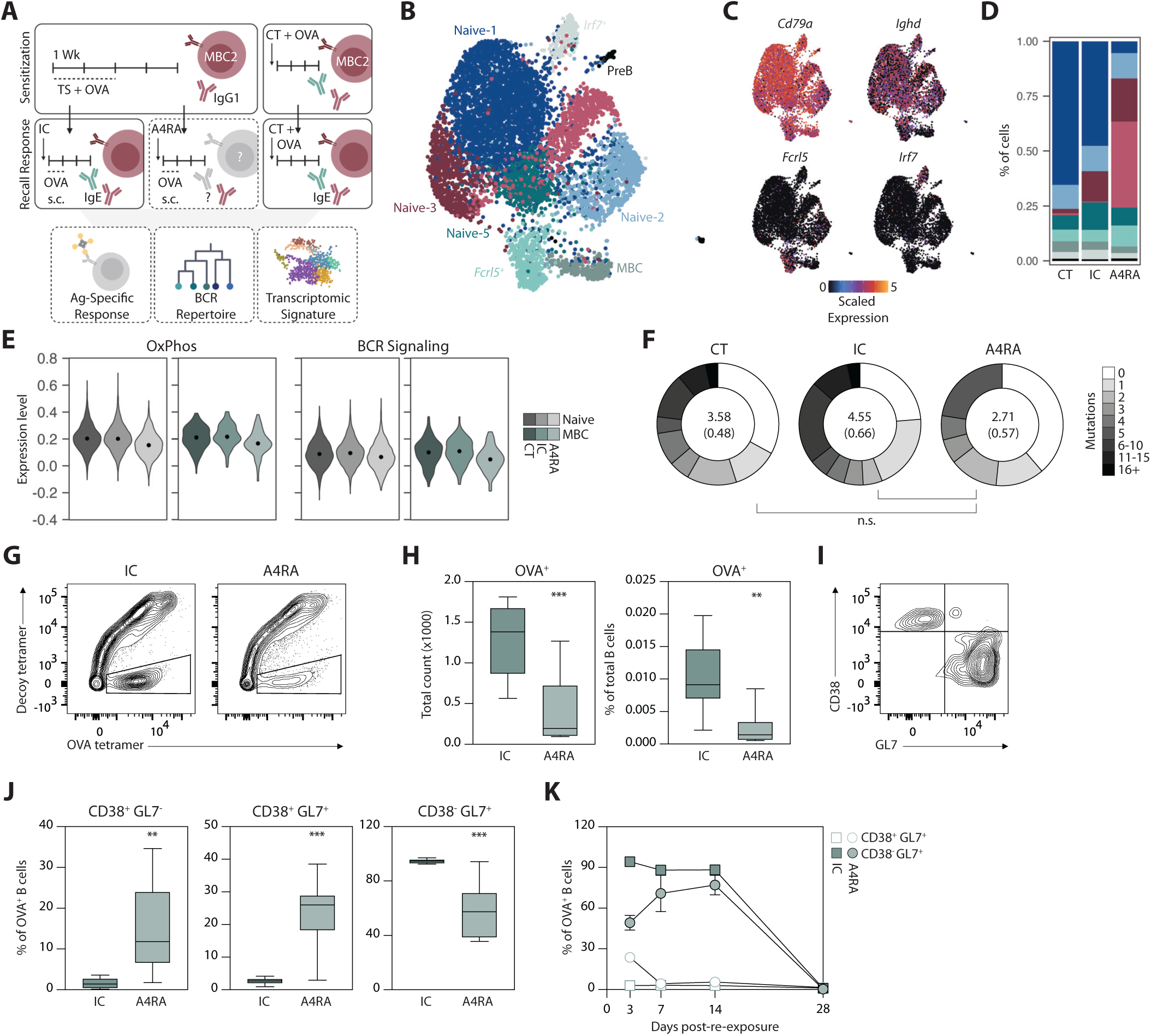
IL-4Ra signaling augments secondary germinal center activity. **(A)** Study schematic. **(B)** UMAP of B cell clusters from scRNA-seq analysis of B cells isolated from spleens and inguinal LNs of OVA-sensitized mice. Mice were either sensitized subcutaneously with CT or epicutaneously by tape-stripping. Epicutaneously sensitized mice were treated with either IC or A4RA prior to s.c. re-exposures and tissue harvest. *n* = 5 CT, *n* = 10 IC, *n* = 10 A4RA. **(C)** UMAP plots of genes defining naïve and MBC clusters. **(D)** Percent of cells in each cluster by treatment group. **(E)** Expression level of gene modules in naïve B cell clusters 1-5 (grey) and MBCs (green) by treatment group. Point indicates median. **(F)** VDJ analysis of MBCs identifying the number of nucleotide mutations compared to the germline sequence. Shading in circle plots depict the fraction of cells with each number of mutations. Mean (SEM) number of mutations are displayed within the circles. **(G)** Representative contour plots of enriched OVA^+^ B cell population from inguinal LNs of epicutaneously sensitized mice. Pre-gated as live > singlet > F4/80^-^ CD3^-^ > B220^+^. **(H)** Summary plots of OVA^+^ B cell total count and frequency. **(I)** Representative contour plot of resting (CD38^+^ GL7^-^), transitional (CD38^+^ GL7^+^) and GC (CD38^-^ GL7^+^) B cells. Pre-gated from OVA^+^ B cells > IgM^-^ IgD^-^. **(J)** Summary plots of OVA^+^ B cell frequency within the populations from (I) at three days post-boost. **(K)** Summary plot of OVA^+^ B cell frequency within transitional (empty) and GC (filled) populations in post-boost time course. Data represent 2 experiments, each with 4 mice per group. Data are presented as median ± min/max (H, J) or mean ± SEM (K). ** *p* < 0.01, *** *p* < 0.001.

Four weeks following immunization and s.c. re-exposures, paired scRNA-seq and scVDJ-seq were performed on 7,369 B cells. Clustering analysis revealed subsets of naïve B cells (*Ighm^+^ Ighd^+^*), class-switched MBCs (*Ighm^-^ Ighd^-^*), and an *Ighd*^+/-^ subset defined by high *Fcrl5* and *Apoe* expression (**Fig. 1B and 1C**). Naïve-1 was the dominant naïve subset in CT and IC-treated mice, though was almost completely replaced by the Naïve-4 cluster in A4RA-treated mice (**Fig. 1D**). Relative to other B cell clusters, Naïve-4 cells exhibited marked transcriptional downregulation, particularly across several metabolic pathways and genes relating to Ag presentation and cellular localization (**fig. S1A and S1B**). Indeed, it has been shown that *in vitro* B cell stimulation with IL-4 initiates metabolic reprogramming and MHC-II upregulation (*22*–*24*). The Naïve-3 subset, which was almost exclusively found in epicutaneously sensitized mice, had upregulated *Il31ra* expression, the ligand (IL-31) of which is implicated in promoting dermatitis (**fig. S1C and S1D**) (*25*). Other naïve and MBC clusters were similarly represented between the three groups. Across all core naïve clusters (1–5) and MBCs, A4RA treatment resulted in downregulated oxidative phosphorylation and B cell receptor (BCR) signaling gene modules – two imperative processes for germinal center (GC) activity (**Fig. 1E**) (*26*, *27*).

Previous reports have demonstrated a critical requirement of IL-4 signaling for primary GC activity in models of type 2 immunity, though it remains unclear whether this requirement is maintained for secondary responses where primary responses can prime MBCs for a GC or PC fate (*28–30*). The requirement of secondary T follicular helper cell-mediated B cell activation to mount an IgE response in epicutaneously sensitized mice was confirmed using CD40L blockade before and during s.c. re-exposures, whereby anti-CD40L prevented OVA-specific IgE production and clinical reactivity (**fig. S2A-C**). To investigate how an IL-4/IL-13 signaling deficiency might impact secondary GCs, we utilized the paired scRNA- and scVDJ-seq data to determine the extent of somatic hypermutation (SHM) within the MBC cluster. Despite an absence of highly mutated sequences with A4RA treatment, the overall extent of SHM was not significantly different between groups (**Fig. 1F**).

As the above findings provided only transcriptional insight within the bulk (non-Ag-specific) B cell repertoire, we assessed the impact of A4RA on OVA-specific B cells by flow cytometry using fluorochrome-conjugated OVA tetramers (*31*, *32*). In agreement with the altered B cell metabolic profile in the transcriptomic data suggestive of compromised B cell survival and/or proliferative capacity, the frequency of OVA^+^ B cells was 3-fold lower in A4RA-treated mice, with an average of 406 versus 1269 cells per dLN pair in A4RA and IC groups, respectively (**Fig. 1G and 1H**). Expression of CD38 and GL7 identifies three populations of B cells differing in their activation states: CD38^+^ GL7^-^ resting naïve/memory (1), CD38^+^ GL7^+^ transitional or multipotent precursor cells (2), and CD38^-^ GL7^+^ GC B cells (3) (**Fig. 1I**) (*33*). At three days post-re-exposures, the A4RA group had significantly impaired OVA-specific B cell activation, evidenced by an increased frequency of CD38^+^ GL7^-^ OVA-specific B cells and a concomitant decrease in the CD38^-^ GL7^+^ population (**Fig. 1J**). However, 23% of OVA-specific B cells in A4RA-treated mice were in a transitional state compared to 2.6% in IC-treated mice, which suggested that loss of IL-4/IL-13 signaling altered the kinetics of MBC reactivation. Indeed, assessment of GC activity at 7-, 14-, and 28-days post-re-exposure demonstrated that the proportion of OVA^+^ B cells that ultimately participate in GCs is no different between groups and A4RA treatment merely stalls early GC formation (**Fig. 1K**). Taken together, these data suggest that loss of IL-4/IL-13 signaling elicits profound transcriptional changes to naïve B cells and MBCs, but GC-dependent recall responses can ultimately proceed.

### MBC2 phenotype is diminished with IL-4Rα blockade in mice and humans

MBC2s have been identified as an important reservoir for pathological IgE responses (*14*, *15*). Previously, IL-4 was shown to be necessary for MBC2 development and, likewise, blockade of IL-4/IL-13 signaling issued a decline in MBC2 frequency (*13*, *14*, *18*). Despite the observed decrease in MBC2s a small number persist, though whether expression of the pathogenic IL-4-responsive phenotype is maintained to the same extent has not been explored.

To address this, we leveraged publicly reposited B cell scRNA-seq datasets involving perturbation of IL-4/IL-13 signaling. Including our own data generated above, our analysis involved four datasets comprising both mouse and human, various models of type 2 immunity, enhanced and abrogated IL-4/IL-13 signaling, and with or without Ag re-exposure (**Fig. 2A**) (*18*, *34*, *35*). For each dataset, unsupervised clustering was performed on B cells (*Cd79a^+^*), which resolved clusters of MBCs classified by absence of *Ighd*, *Tcl1a* (human), and *Bcl6* (mouse) and the presence of *Hhex* (mouse) and *Ccr6* (mouse) expression (**Fig. 2B and S3A-D**). *Fcer2*, *Il4r*, and *Ighg1* expression was compared between each control and interventional group to evaluate the impact of enhanced or inhibited IL-4/IL-13 signaling on MBC2-related gene expression (**Fig. 2C**). *Fcer2* expression in MBCs was consistently downregulated across datasets involving IL-4/IL-13 signaling inhibition. IL-4Rα blockade also elicited a decline in *Il4r* expression, apart from epicutaneously sensitized mice where *Il4r* transcript abundance was also low in IC mice. Conversely, with increased IL-4/IL-13 bioavailability to B cells through elimination of its uptake by lymphoid stromal cells (*Ccl19^Cre/+^ Il4ra^fl/fl^* bone marrow chimeras; LSC Il4r deficiency), *Fcer2* and *Il4r* expression was enhanced within the MBC cluster (*34*). Intriguingly, *Ighg1* expression was downregulated only in experimental systems involving Ag re-exposure during IL-4Rα blockade.

**Fig. 2.**
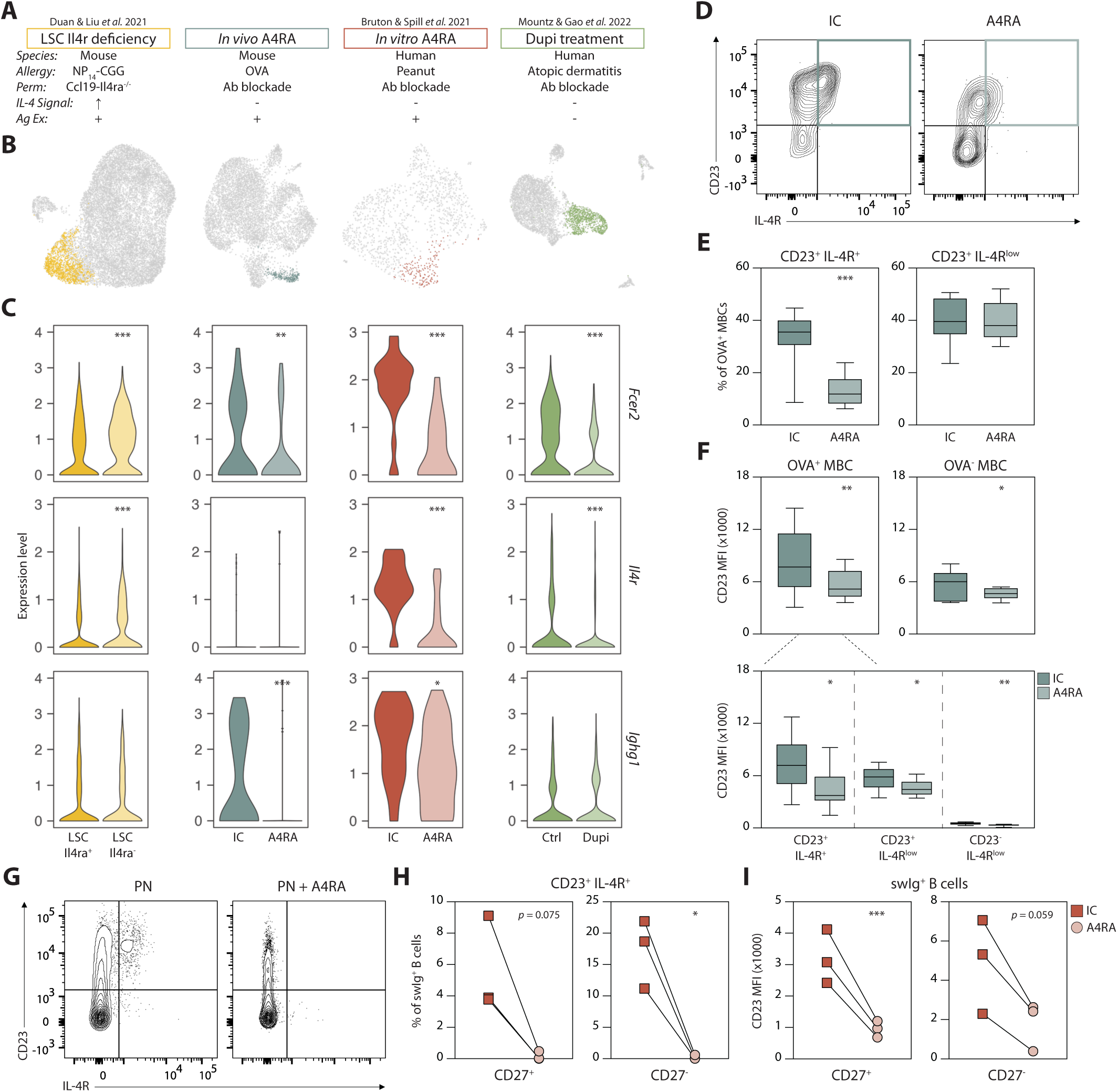
MBC2 phenotype is ameliorated with IL-4Ra blockade in mice and humans. **(A)** Overview of analyzed datasets. **(B)** UMAP of B cell clusters from scRNA-seq datasets. Colored points indicate class-switched cluster used for downstream analyses. **(C)** Expression level of MBC2-related genes in MBC clusters. **(D)** Representative contour plots of OVA^+^ MBC2s from inguinal LNs of epicutaneously sensitized mice four weeks after re-exposure and Ab treatment. Pre-gated as OVA^+^ B cells > CD38^+^ GL7^-^ > IgM^-^ IgD^-^. **(E)** Summary plots of MBC2 and CD23^+^ IL-4R^low^ MBC frequency from IC and A4RA-treated epicutaneously sensitized mice. **(F)** MFI of CD23 within OVA^+^ (top left) and OVA^-^ (top right) MBCs from IC and A4RA-treated epicutaneously sensitized mice and faceted by IL-4R and CD23 expression (bottom) within OVA^+^ population. Pre-gated as OVA^+/-^ B cells > CD38^+^ GL7^-^ > IgM^-^ IgD^-^. **(G)** Representative contour plots of MBC2s from human PBMCs 8 days following *in vitro* allergen stimulation and treatment with IC or A4RA. Pre-gated as live > singlet > CD3^-^ CD14^-^ CD16^-^ CD19^+^ > IgD^-^ IgM^-^ > CD38^low-med^. **(H)** Summary plots of human MBC2 frequency in **G**, separated by CD27 expression. **(I)** MFI of CD23 within CD27^+/-^ class-switched B cells from human PBMCs 8 days following *in vitro* allergen stimulation and treatment with IC or A4RA. Data represent 2-3 experiments with 5 mice per group and are presented as median ± min/max (E, F) or 2 experiments (*n* = 3 blood donors) with each line representing a single donor (H, I). * *p* < 0.05, ** *p* < 0.01, *** *p* < 0.001. LSC, lymphoid stromal cell; Perm, permutation; Re-Ex, re-exposure.

To corroborate the diminished MBC2 phenotype within Ag-specific cells, we assessed co-expression of surface IL-4Rα and CD23 on tetramer^+^ MBCs by flow cytometry four weeks following *s.c.* re-exposures in epicutaneously sensitized mice (**Fig. 2D**). A4RA-treated mice exhibited a 61% reduction in the frequency of IL-4Rα^+^ CD23^+^ cells in the OVA-specific MBC compartment (**Fig. 2E**). IL-4Rα did not appear to be maintained on the cell surface bound to A4RA, as surface rat-IgG2a (A4RA Fc region) was undetectable at this timepoint (**fig. S4**). Consistent with a role of IL-4 in inducing CD23 expression, A4RA treatment decreased surface expression of CD23, even amongst IL-4R^low^ cells (**Fig. 2F**) (*36*). CD23 downregulation was not, however, unique to Ag-specific cells, but instead was broadly observed throughout non-specific MBCs (**Fig. 2F**).

To also validate the human scRNA-seq observations, we stimulated peripheral blood mononuclear cells (PBMCs) from peanut-allergic donors (*n*=3) with crude peanut extract for eight days in the presence or absence of A4RA. We have previously shown that 8% of MBCs in unstimulated cultures exhibit an MBC2-like transcriptional profile and expand to over 20% upon peanut stimulation. Extending upon this to elucidate how A4RA treatment impacted protein expression of MBC2 markers, we could observe CD23^+^ IL-4Rα^+^ class-switched B cells in stimulated cells but these were virtually undetectable with A4RA treatment (**Fig. 2G**). Consistent with previous reports, CD23^+^ IL-4Rα^+^ cells had variable CD27 expression, though we found that both populations were similarly impacted by A4RA (**Fig. 2H**) (*13*, *14*). Moreover, as was observed in mice, A4RA broadly decreased CD23 mean fluorescence intensity (MFI) within class-switched B cells (**Fig. 2I**). Collectively, these data demonstrate a conserved requirement for IL-4/IL-13 signaling to maintain heightened type 2 polarization within established MBC compartments, which contests a previous suggestion regarding MBC2 phenotypic resiliency (*13*). These data also raise the issue of how Ag re-exposure during IL-4Rα blockade may skew the isotype distribution away from IgG1 – the dominant intermediary for IgE CSR.

We acknowledge that, despite using different Ab clones for IL-4Rα blockade and detection by flow cytometry, it is possible that steric hindrance may have diminished the detection of surface IL-4Rα bound by the blocking Ab. Indeed, engagement of surface IL-4Rα with dupilumab (anti-human IL-4Rα blocking Ab) promotes IL-4Rα internalization via endocytosis, although a dupilumab-bound fraction is also retained on the cell surface (*37*). Nevertheless, the absolute amount of IL-4Rα (both free and blocking Ab-bound) decreases in dupilumab-treated patients, supporting the observed reduction in IL-4Rα expression on OVA^+^ MBCs in our study (*37*, *38*). Moreover, our transcriptomic analyses indicate a diminished MBC2 phenotype, as both *Fcer2* and *Il4r* expression was significantly decreased in human MBCs that received A4RA treatment.

### Long-lived Ag-specific IgG2b/c responses emerge in the absence of IL-4/IL-13 signaling

As we observed that recall responses occurred in the absence of IL-4/IL-13 signaling, alongside dampened MBC2 polarization (including *Ighg1*), we sought to define the Ig product of recall responses. Evaluating isotypes downstream to IgG1 – the dominant memory reservoir in epicutaneously sensitized mice – an increase in *Ighg2b* and *Ighg2c* transcripts was observed in the MBC cluster of A4RA-treated mice (**Fig. 3A**). *Igha* transcript expression was also increased in A4RA-treated mice, though not significantly. Consistent with the notion that IgE^+^ MBCs are exceptionally rare or non-existent in mice, mature *Ighe* transcripts were not readily detectable in any experimental group (6 of 7369 cells) (*39–41*). From the paired scVDJ-seq analysis we plotted the assigned *Ighc* gene for each MBC with at least three nucleotide mutations, suggestive of hypermutated, Ag-experienced cells (**Fig. 3B**). The distribution of isotype usage in mutated MBCs was comparable between CT and IC mice, except for *Ighg2b*, which had a substantially greater presence in CT in agreement with the reported CT-induced secretion of TGF-β, a driver of IgG2b CSR (*42*, *43*). By contrast, A4RA induced a pronounced increase in *Ighg2c* above both CT and IC – an isotype that is not associated with type 2 immunity.

**Fig. 3.**
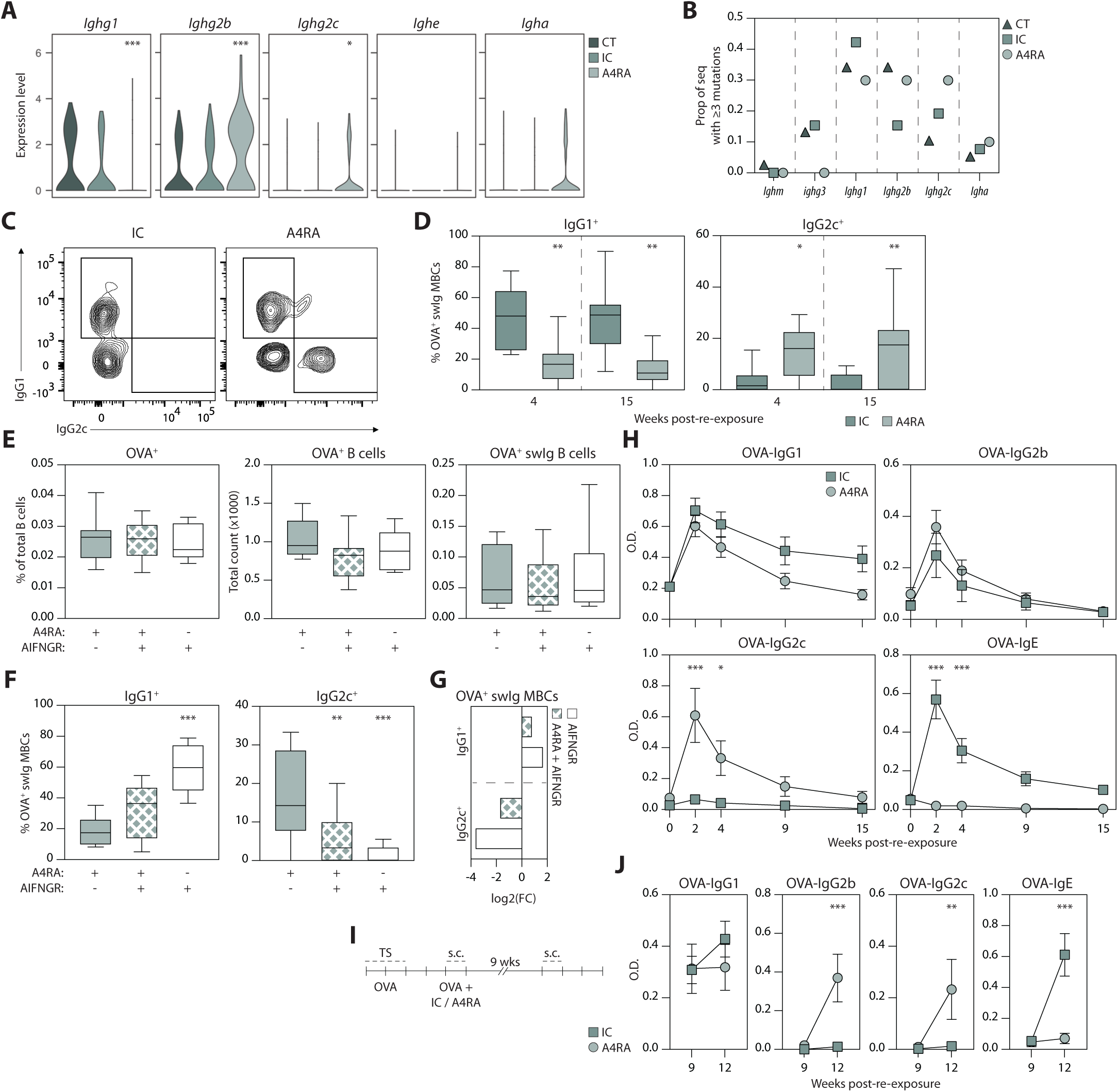
Longitudinal analysis of Ag-specific B cell responses with absent IL-4 signaling. **(A)** Expression level of *Ighc* genes inclusive and downstream of *Ighg1* from scRNA-seq analysis of B cells isolated from spleens and inguinal LNs of CT-immunized mice and A4RA/IC-treated epicutaneously sensitized mice. Gene expression is plotted for cells within the MBC cluster (Fig. 1B). **(B)** VDJ analysis of MBC cluster. Plot depicts the assigned *Ighc* gene in MBCs with ≥3 nucleotide mutations compared to the germline sequence. **(C)** Representative contour plots of IgG1^+^ and IgG2c^+^ populations from inguinal LNs of IC/A4RA-treated epicutaneously sensitized mice four weeks post-boost and Ab treatment. **(D)** IC/A4RA-treated mice were rested up to 15 weeks after re-exposures. IgG1^+^ (left) and IgG2c^+^ (right) MBC frequency are plotted at four and 15 weeks post-boost. Pre-gated as OVA^+^ B cells > CD38^+^ GL7^-^ > IgM^-^ IgD^-^. **(E)** Epicutaneously sensitized mice were treated with A4RA, AIFNGR, or both at the time of subcutaneous re-exposure. Summary plots of OVA^+^ B cell and swIg OVA^+^ B cell total count and proportion of total B cells. **(F)** IgG1^+^ (left) and IgG2c^+^ (right) MBC frequency are plotted at four weeks post-boost. Pre-gated as OVA^+^ B cells > CD38^+^ GL7^-^ > IgM^-^ IgD^-^. **(H)** OVA-specific serum Ig profiles of IC/A4RA-treated mice at zero to 15 weeks post-boost. **(I)** IC/A4RA-treated mice were rested for nine weeks to allow for clearance of OVA-specific IgG2c and clearance of IC/A4RA blocking antibodies. Subsequently, mice received tertiary OVA exposures and serum was collected at week 12. **(J)** OVA-specific serum Ig profiles are depicted before and after tertiary exposures. (C-J) Data represent 2 experiments with at least 4 mice per group. Data are presented as mean ± SEM (H, J), median ± min/max (D-F), and mean (G). * *p* < 0.05, ** *p* < 0.01, *** *p* < 0.001. Seq, sequences; swIg, class-switched; AIFNGR, anti-IFN-gR1; OD, optical density; Wks, weeks.

To assess whether the skewed isotype usage occurred within the Ag-specific B cell repertoire and the longevity of such response, OVA^+^ MBCs and the OVA-specific serum Ig profile were interrogated up to 15 weeks after the recall response. OVA^+^ IgG1^+^ MBCs were present in both IC and A4RA-treated mice, though with a significantly decreased frequency with A4RA treatment (**Fig. 3C and 3D**). By comparison, OVA^+^ IgG2c^+^ MBCs emerged with A4RA treatment and were sustained beyond three months. As IgG2c CSR is driven by IFN-γ and our previous work demonstrated an increase in IFN-γ production with A4RA treatment, we sought to establish the role of IFN-γ in the emergence of OVA^+^ IgG2c^+^ MBCs (*18*, *44*, *45*). In mice treated with A4RA, A4RA with concurrent IFN-γ receptor 1 blockade (AIFNGR), or AIFNGR alone, the frequency of OVA^+^ B cells was similar, suggesting that IFN-γ signaling does not influence Ag-specific B cell expansion in a recall response (**Fig. 3E**). The emergence of OVA^+^ IgG2c^+^ MBCs, however, was halted with concurrent AIFNGR (**Fig. 3F and 3G**).

Skewed isotype usage in the absence of IL-4 signaling was also reflected in the analysis of serum OVA-specific Igs (**Fig. 3H**). As per our previous observations, serum OVA-IgE responses were short-lived in IC mice with a completely abrogated response in A4RA-treated mice (*18*, *21*). Consistent with the induction of IgG2c^+^ MBCs, serum OVA-IgG2c was observed with A4RA treatment though was not long-lived and declined at a similar rate to IgE in IC mice. Disparate to the reduction in OVA^+^ IgG1^+^ MBCs, Ag re-exposure induced a comparable resurgence of serum OVA-IgG1 in both groups, suggesting that A4RA may have limited impact on MBCs fated for PC differentiation. Serum OVA-IgG2b titers were comparable between both groups and OVA-IgA was undetectable.

We reasoned that if the skewed isotype response by A4RA was stably maintained then a resurgence of IgG2c should be facilitated by seeded IgG2c^+^ MBCs. To this end, mice were provided tertiary OVA re-exposures nine weeks post-A4RA/IC and s.c. OVA administration (**Fig. 3I**). At this point there is clearance of both serum OVA-IgG2c and the blocking A4RA Ab (*18*). Indeed, tertiary Ag re-exposure provided a significant regeneration of OVA-IgG2b and OVA-IgG2c though abrogation of OVA-IgE production remained (**Fig. 3J**). Collectively, these data demonstrate that induction of a recall response with IL-4Rα blockade permit the emergence of a non-type 2 Ab response and propose that this reprogrammed B cell response may be maintained even following A4RA clearance.

LNs at distinct anatomical sites have been shown to differ in their predisposition to promote inflammatory or tolerogenic responses towards Ags (*46*). In this regard, we also investigated A4RA-induced isotype skewing in the context of oral immunization, which preferentially issues tolerogenic IgA-dominant responses. As previously shown, one intragastric administration of Ag + CT does not issue a humoral response (*47*, *48*), though Ag-specific Igs arise after subsequent oral administration of Ag alone (*47*). We adopted this framework to provide systemic administration of IC or A4RA prior to each oral Ag re-exposure (**fig. S5A**). Expectedly, IC-treated mice exhibited a classical type 2 response marked by OVA^+^ IgG1^+^ MBCs in gut dLNs and serum OVA-IgG1 and -IgE (**fig. S5B and S5C**). Mirroring observations from epicutaneous sensitization, A4RA elicited the emergence of OVA^+^ IgG2c^+^ MBCs and serum OVA-IgG2b and -IgG2c, while preventing OVA-IgE synthesis. Conversely, OVA re-exposures did initiate an IgA response which was abrogated with A4RA, reconciling the requirement of IL-4 signaling for mucosal IgA responses (**fig. S5C**) (*49*). Thus, loss of IL-4/IL-13 signaling during Ag re-exposure preferentially promotes IgG2b/c responses irrespective of tissue site.

### Type 1-polarizing adjuvant can redirect isotypic fate of class-switched MBCs arising in type 2-polarizing conditions

Having demonstrated that IFN-γ supports the reprogrammed fate of OVA^+^ MBCs established in type 2-polarizing conditions, we next assessed whether use of a type 1-polarizing adjuvant could also redirect class-switched MBCs to IgG2b/c.To this end, we utilized an adjuvanted re-exposure system to assess whether the type 1 adjuvant, CpG, can divert the Ag-specific response in type 2 immunized mice (**Fig. 4A**). Similar to the effects of A4RA and comparable to that observed in type 1 immunized mice, re-exposure with CpG facilitated an increased frequency of OVA^+^ IgG2c^+^ MBCs and a decline in MBC2s, though the heightened frequency of OVA^+^ IgG1^+^ MBCs remained (**Fig. 4B and 4C**). The emergence of IgG2c^+^ MBCs was also reflected in increased serum titers of OVA-IgG2c, along with OVA-IgG2b (**Fig. 4D**). Remarkably, the type 1-adjuvanted re-exposure did not prevent OVA-IgE production, suggesting that this approach has an attenuated potency in redirecting pathogenic IgE responses relative to A4RA treatment.

**Fig 4.**
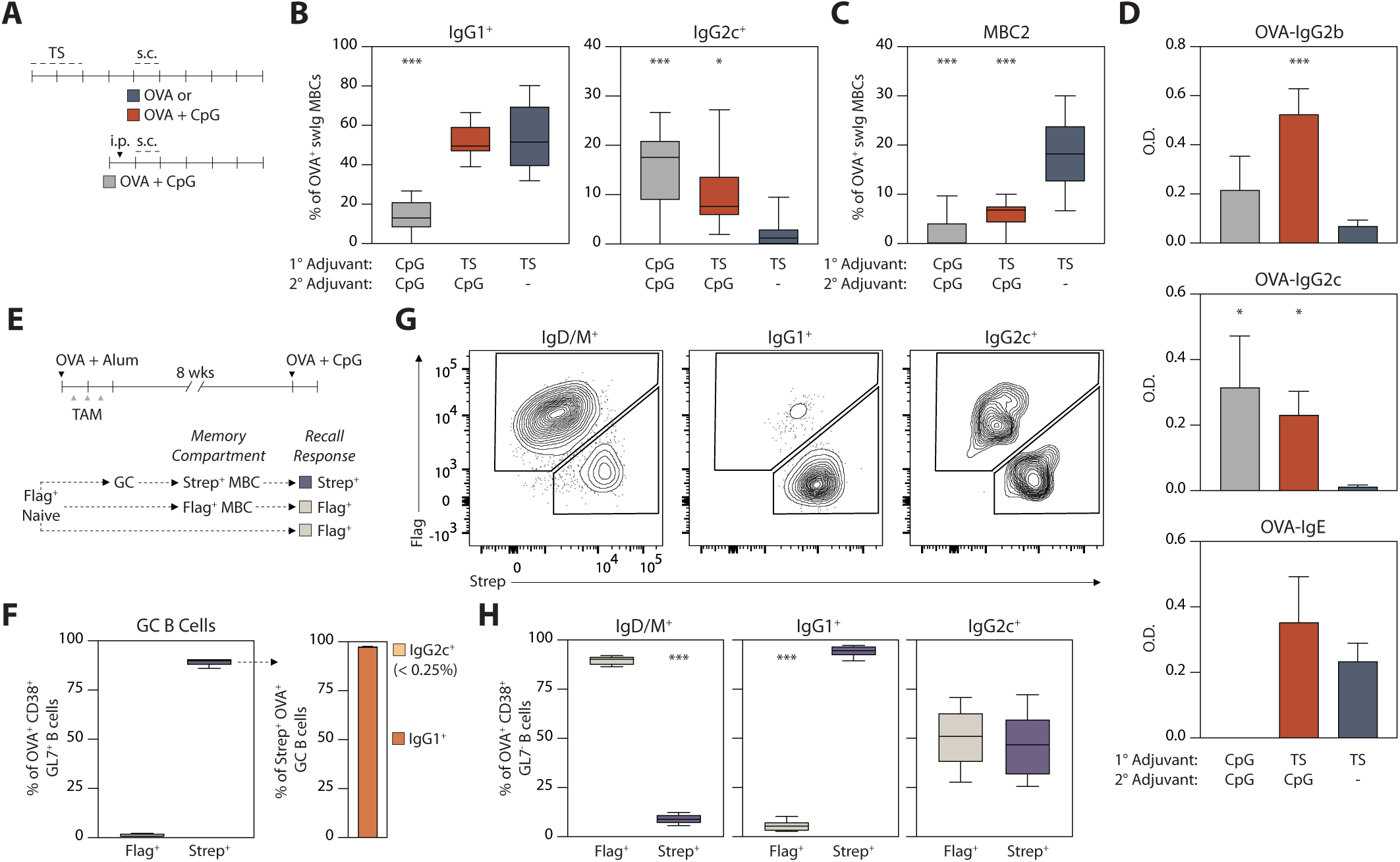
Boosting with type 1 adjuvant reprograms the type 2-primed response. **(A)** Experimental schematic. **(B,C)** IgG1^+^ (left) and IgG2c^+^ (right) MBC frequency (B) and MBC2 cell frequency (C) are plotted at four weeks post-secondary exposures with OVA alone or OVA + CpG. Pre-gated as OVA^+^ B cells > CD38^+^ GL7^-^ > IgM^-^ IgD^-^. **(D)** OVA-specific serum Ig profiles. **(E)** Experimental schematic utilizing *S1pr2-Igk^Tag^* mice. **(F)** Summary plots of Flag/Strep expression in OVA^+^ CD38^-^ GL7^+^ B cells at 7 days post-TAM treatment (left) and IgG1/IgG2c expression in the Strep^+^ population (right). **(G)** Representative contour plots of Flag/Strep expression in OVA^+^ CD38^+^ GL7^-^ B cells from inguinal LNs one week after the recall response with OVA + CpG. **(H)** Summary plots of Flag/Strep expression in IgD/M^+^ (left), IgG1^+^ (middle), and IgG2c^+^ (right) populations. Pre-gated as OVA^+^ B cells > CD38^+^ GL7^-^. Data represent 2 experiments with at least 4 mice per group. Data are presented as mean ± SEM (D) or median ± min/max (B, C, F and H). * *p* < 0.05, ** *p* < 0.01, *** *p* < 0.001. OD, optical density; TAM, tamoxifen; Wks, weeks.

While A4RA provides direct blockade of IL-4/IL-13 signaling in B cells and T cells, the effect of CpG on CD4 T cells likely operates primarily through an indirect mechanism, involving the modulation of Ag-presenting cells (*50*, *51*). Therefore, we hypothesized that a CpG adjuvanted re-exposure may be more effective in ameliorating the pathogenic IgE fate of MBCs when not having to contend with the existing type 2 bias in Ag-experienced CD4 T cells. To test this, we utilized an adoptive transfer approach in which class-switched OVA-enriched MBCs were sorted from OVA + Alum-immunized CD45.1^+^ donor mice and were transferred to B cell-deficient (muMT) CD45.2^+^ naïve mice (**fig. S6A and S6D**). Recipient mice were then immunized with either OVA + CT or OVA + CpG. Of note, OVA + Alum immunization yields an IgG1 and IgE dominant response, where over 85% of the class-switched MBC pool is IgG1^+^ and less than 4% are IgG2b/c^+^, with greater than 50% of the IgG1^+^ also harboring the MBC2 phenotype (*52*) (**fig. S6B and S6C**). Thus, any emergence of OVA-IgG2 responses in recipient mice is not likely to represent an expansion of donor IgG2 clones. Moreover, OVA + CT immunization produces more OVA^+^ IgG2b/c^+^ MBCs than Alum; therefore, if any IgG2 clones in the donor pool do contribute to the OVA-IgG2 response in recipients, we would expect to observe this with both CT and CpG immunization (**fig. S6B**).

One week following adoptive transfer and immunization, donor (CD45.1^+^) OVA^+^ B cells and OVA-specific Igs were detected in recipient mice. As anticipated, immunization of recipient mice with OVA + CT drove the emergence of OVA^+^ IgG1^+^ MBCs and serum OVA-IgG1 and - IgE. In contrast, OVA^+^ IgG2c^+^ MBCs and serum OVA-IgG2b and -IgG2c became present in recipient mice immunized with OVA + CpG, with a sustained absence of OVA-IgE (**fig. S6E and S6F**). Collectively, these data support that Ag re-exposure in the presence of a type 1 adjuvant can redirect the fate of class-switched MBCs established under type 2-polarizing conditions.

### IgG2c^+^ B cells arising in a type 1-adjuvanted recall response emerge from GC-derived and GC-independent cellular origins

MBCs, including MBC2s, can be generated both extrafollicularly (EF) and as an output of GCs (*14*, *53*). To determine whether EF- or GC-dervied MBCs differentially contribute to the reprogrammed IgE response, we utilized a molecular fate mapping approach with *S1pr2-Igk^Tag^* mice (*54*). These mice harbor a LoxP-flanked Flag tag and downstream Strep tag at the Igκ light chain gene and Cre^ERT2^ expression under control of the GC-specific *S1pr2* gene (*54*). In this model, B cells and their secreted antibodies express a Flag tag, unless engaged in GCs during tamoxifen (TAM) treatment, at which point the Flag tag is irreversibly replaced with a Strep tag.

Following previously established methodology, TAM was administered to *S1pr2-Igk^Tag^* mice beginning four days after OVA + Alum immunization during GC activity (**Fig. 4E**) (*54*). The product of this primary response yields class-switched cells that are Strep^+^ if they were GC-derived or Flag^+^ arising from an EF pathway. Seven days following the final TAM treatment, 89.4% of GC B cells were Strep^+^ (**Fig. 4F**). Of Strep^+^ GC B cells, 97.4% were IgG1^+^ and less than 0.25% were IgG2c^+^, which is in agreement with robust GC activity and IgG1 CSR known to arise in Alum immunization models (*55*). Two months later, following contraction of GC activity, mice were re-exposed to OVA + CpG (without additional TAM treatment) and the OVA^+^ B cell compartment was analyzed by flow cytometry (**Fig. 4G**). As expected, the overwhelming majority of OVA^+^ B cells expressing IgD/IgM were Flag^+^ whereas IgG1^+^ cells were mostly Strep^+^, indicative of their GC-derived origin (**Fig. 4G**). OVA^+^ IgG2c^+^ B cells, however, exhibited a equal distribution of Flag and Strep expression, suggesting that the IgG2c^+^ B cells emerging in the type 1-adjuvanted recall response can originate from both naïve B cells or EF-derived MBCs (Flag^+^) and GC-dervied MBCs (Strep^+^). Given that the product of Alum immunization was overwhelming IgG1-dominant, it is likely that Strep^+^ IgG2c^+^ B cells arose from GC-derived IgG1^+^ MBCs (**fig. S5B**). In this regard, we next tested this possibility by assessing the VDJ repertoire and clonal origins of OVA-specific B cells.

### Class switch origins of IgG2b/c^+^ clones are comprised of multiple isotypes

The observed loss of MBC2 polarization and divergence towards non-IgE humoral response indicate the potential to reprogram pathogenic IgE responses; however, it remains unclear whether skewed isotype usage arises *de novo* (i.e., from naïve or unswitched B cells) or from direct reprogramming of existing class-switched MBCs. To address this, we first looked for the presence of germline sterile transcripts (S-Tx), the expression of which precedes CSR to the productive Ig isotype (*56*, *57*). Thus, the presence of *Ighg2b/c* S-Tx in cells with productive IgM, IgG3, or IgG1 would indicate B cells that are likely primed for IgG2b/c CSR. We used the sciCSR pipeline to annotate S-Tx expression in our scRNA-seq dataset (**Fig. 5A**). S-Tx inclusive of and downstream to *Ighg1* were preferentially expressed in the MBC cluster, congruent with cells in this cluster being Ag-experienced (**Fig. 5B**). *Ighg3* S-Tx were most prevalent in the *Fcrl5^+^* cluster which aligns with the observed IgG3 bias in atypical (*Fcrl5^+^*) MBCs (*58*, *59*). *Ighg2b* S-Tx were also present in naïve-like clusters, perhaps identifying recently activated B cells yet to undergo CSR from IgM.

**Fig. 5.**
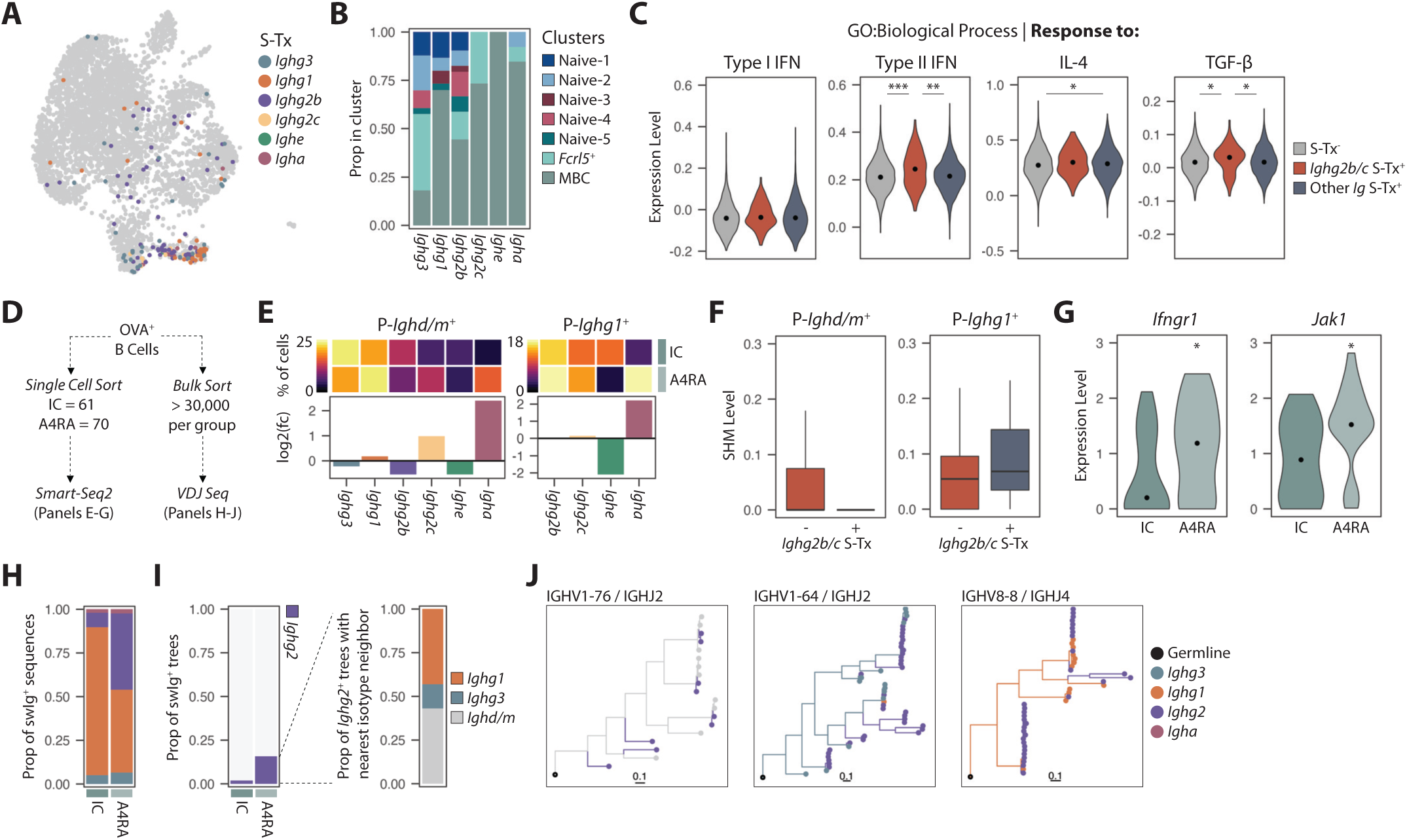
Clonal analysis of polyclonal and OVA-specific B cells. **(A)** UMAP of B cell clusters from scRNA-seq (Fig. 1B). Colored points indicate the most downstream *Igh* S-Tx expressed by a cell. **(B)** Distribution of S-Tx amongst B cell clusters. **(C)** Expression level of gene modules in B cells without S-Tx (grey), B cells with *Ighg2b/c* S-Tx (red), and B cells with other S-Tx (blue). Point indicates median. **(D)** Schematic depicting use of OVA-specific B cells for Smart-Seq2 and bulk VDJ sequencing experiments. **(E)** Heatmap displaying the percent of B cells expressing S-Tx downstream to their productive isotype and barplots depicting the fold change in S-Tx (IgD/M, left; IgG1, right) with A4RA treatment compared to IC. **(F)** SHM frequency between *Ighg2b/c* S-Tx^+^ (red) and *Ighg2b/c* S-Tx^-^ (blue) B cells with productive IgD/M (left) or productive IgG1 (right). Data are presented as median ± min/max. **(G)** Expression level of genes from Smart-Seq2 analysis. Gene expression is plotted for cells with a productive *Ighg1* call from MBC clusters (*n*=22 cells per group). **(H)** Isotype usage amongst swIg sequences. **(I)** Proportion of clonal lineage trees containing *Ighg2* amongst trees with ≥ 2 isotypes (left) and nearest isotype neighbor amongst *Ighg2*-containing trees (right). **(J)** Representative clonal lineage trees containing *Ighg2* sequences and upstream isotypes (*Ighd/m*, *Ighg3*, and *Ighg1*) from A4RA-treated mice. Scale indicates IGHV SHM frequency. * *p* < 0.05, ** *p* < 0.01, *** *p* < 0.001. P, productive; Prop, proportion.

We then asked whether B cells with *Ighg2b/c* S-Tx differed in their gene expression profile relating to cytokine signaling involved in isotype determination during CSR. We found that *Ighg2b/c* S-Tx^+^ cells had significantly higher expression of genes related to responsiveness to type II IFN and TGF-β, but not type I IFN or IL-4 (**Fig. 5C**). Conversely, cells expressing S-Tx other than *Ighg2b/c* had heightened expression of genes involved in the response to IL-4 relative to S-Tx^-^ cells.

Given that these data indicated a potential for IgG2b/c CSR from naïve and class-switched polyclonal cells, we posited that this would also be true of OVA-specific cells. Accordingly, we next performed scRNA-seq (Smart-Seq2) and bulk VDJ sequencing of OVA-specific B cells sorted from IC- and A4RA-treated mice (**Fig. 5D**). Smart-Seq2 yielded the reconstruction of full length BCRs for 61 and 70 OVA^+^ B cells from IC- and A4RA-treated mice, respectively. Of the sequenced cells, 70 were class-switched. We employed the same approach described above to investigate the presence of S-Tx in OVA-specific B cells with a productive IgD/M call or IgG1 call. With A4RA treatment there was a near complete absence of *Ighe* S-Tx in IgG1^+^ OVA- specific B cells (2.6% vs. 11.5% of IgG1^+^ in A4RA and IC, respectively), confirming deterred IgE CSR (**Fig. 5E**). A4RA-treated cells also exhibited a 2-fold increase in OVA^+^ IgD/M^+^ B cells expressing *Ighg2c* S-Tx, though the proportion of OVA^+^ IgG1^+^ B cells with *Ighg2c* S-Tx was comparable between groups (**Fig. 5E**). The presence *Ighg2b/c* S-Tx was unrelated to the level of SHM in IgD/M^+^ and IgG1^+^ cells, suggesting that this is not reflective of increased Ag exposure (**Fig. 5F**). We also further explored the dataset from LSC Il4r-deficiency model and found that the reverse held true – with increased IL-4 bioavailibity, MBCs harbored less *Ighg2b* and *Ighg2c* S-Tx, and increased *Ighe* S-Tx, corroborating that diversion to an IgG2b/c trajectory requires suppressed IL-4 signaling in B cells (**fig. S7A-C**). Notably, A4RA treatment heightened the proportion of both IgD/M^+^ and IgG1^+^ cells with *Igha* S-Tx, with increases of 5.4-fold and 4.8-fold, respectively. As there was not a detectable increase in serum OVA-IgA, the increase in *Igha* S-Tx may signal a later propensity for IgA CSR with continued Ag re-exposures.

Finally, amongst MBCs with productive *Ighg1*, we compared the expression of genes comprising the IFN-γ receptor heterodimer and their upstream signaling kinases, given our finding that IFN-γ is necessary for the A4RA-induced IgG2c response (**Fig. 3F**). While no difference was observed for *Ifngr2* and its associated signaling kinase *Jak2*, *Ifngr1* and its signaling kinase *Jak1* were significantly upregulated in OVA^+^ *Ighg1*^+^ MBCs from A4RA-treated mice and were downregulated in MBCs from the LSC Il4r-deficiency model (**Fig. 5G and fig. S7D**). Of relevance, *Ifngr2* is constituvely expressed at low levels, whereas *Ifngr1* expression can be upregulated by pro-inflammatory cytokines, including IFN-γ itself, which coincides with increased IFN-γ production observed in A4RA-treated mice (*60*, *61*).

The limited throughput of the Smart-Seq2 approach did not permit the detection of clonally related sequences within either treatment group. In this regard, we utilized bulk VDJ sequencing of OVA-specific B cells sorted from pooled LNs and spleens to assess the clonal ancestry of *Ighg2b/c*^+^ cells. Parallel analyses with Ig sequences derived in the 10X dataset were carried out to gain insight as to whether the effects of A4RA are occurring broadly throughout all B cells or are contained within the Ag-specific repertoire. IL-4/IL-13 signaling did not influence CDR3 length in unswitched and class-switched B cell repertoires (**fig. S8A and S8B**). Amongst OVA-specific B cells, the extent of SHM was higher in *Ighg1* and *Ighg2b/c* sequences from A4RA-treated mice, which diverges from observations in bulk B cells (**fig. S8C and S8D**). IL-4 signaling has been shown to modulate BCL6 expression, leading to the output of GC-derived MBCs (*62*). Therefore, it is possible that blockade of IL-4 signaling permitted sustained GC activity enabling further rounds of SHM in OVA-specific cells. We also observed apparent biases in V gene usage within the class-switched repertoire of OVA-specific B cells (**fig. S8E**); however, this was also present within polyclonal B cells and thus is unlikely related to the OVA-specific recall response (**fig. S8F**). Regarding isotype usage, over 40% of class-switched sequences in the A4RA group were *Ighg2b/c*, compared to only 8% in IC, which coincidences with that observed at the transcriptional, cellular, and protein level (**Fig. 5H**).

Broadly probing whether unswitched or class-switched cells were putative precursors to IgG2b/c clones, we first measured the correlation of paired IGHV and IGHJ gene usage between isotypes. Expectedly, paired IGHV-IGHJ gene usage was most highly correlated between *Ighm* and *Ighd* clones in both groups (**fig. S8G**). IGHV-IGHJ gene usage in *Ighg2b/c* clones also correlated most strongly with *Ighm* and *Ighd* clones, and was significantly correlated with *Ighg3* and *Ighg1* clones in A4RA but not IC. Next, we partitioned sequences into clonal lineages through hierarchical clustering, as described previously (*63*, *64*). Amongst trees containing class-switched sequences, 16% of trees in A4RA contained *Ighg2b/c,* versus 2% in IC (**Fig. 5I**). Furthermore, within the *Ighg2b/c*-containing trees in the A4RA group, 57% also included upstream class-switched sequences (*Ighg3* and/or *Ighg1*), confirming a shared clonal lineage between *Ighg2b/c* and other class-switched cells. Accordingly, mutated *Ighg2b/c* sequences were present in B cell lineage trees containing expanded IgM, IgG3, and IgG1 clones (**Fig. 5J**). Collectively, *Ighg2b/c*^+^ cells exhibited clonal relationships with all upstream isotypes, demonstrating that both naïve and class-switched OVA-specific B cells can give rise to IgG2b/c *in vivo*.

## DISCUSSION

MBCs, compared to naïve counterparts, acquire enhanced functional and metabolic properties increasing fitness for recall responses to the same Ag. The MBCs established upon allergic sensitization gain a unique propensity to switch to IgE upon subsequent allergen exposures, under the influence of IL-4. Whether this specialized B cell memory response is functionally plastic was not yet established. Assessing transcriptional profile, BCR repertoire, Ag-specific cellular memory, and humoral Ab responses, our work identifies the capacity to reprogram the pathogenic IgE-fated response towards a type 1-like, non-pathogenic IgG2c program. Numerous studies in mice and humans have now established that IgG1 is the dominant MBC reservoir for IgE (*5*, *10*, *11*, *21*, *65*). Recent work by us and others has extended upon this concept illustrating that IgE-fated MBCs (“MBC2s”) express ε germline transcripts and upregulate surface expression of CD23 and IL-4Rα, seemingly remaining prepared for IL-4 responsiveness (*13–15*). In the absence of IL-4/IL-13 signaling in a recall response, we have previously shown that IgE responses are indeed averted, and cytokine production skews toward a Th1 profile (*18*). In the present work investigating how IL-4Rα blockade impacts the B cell repertoire beyond avoidance of IgE production, we found that loss of IL-4/IL-13 signaling *in vivo* broadly suppresses gene expression, particularly relating to metabolic processes and BCR signaling. Intriguingly, Ag re-exposure could still elicit Ag-specific GC activity in the absence of IL-4/IL-13, an issue which has remained contentious (*28*, *62*, *66*).

As a primary product of a GC response is PC production, we were particularly intrigued by the sustained absence of serum IgE without a compensatory increase in IgG1. Coinciding with our previous observations of an increased IFN-γ to IL-4 ratio, we found that A4RA treatment induced an IFN-γ-dependent Ag-specific IgG2c response, which was otherwise absent with IC. The effector functions of IgG2c are primarily achieved through FcγRI and FcγRIV expressed on myeloid cells (*67*). Engagement with these receptors facilitates effector functions, such as clearance of immune complexes and Ab-dependent cell cytotoxicity, thereby playing an important role in protection against pathogens and a notable absence in type I hypersensitivities. The functional repercussions of an increase in allergen-specific Ab isotypes associated with type 1 immunity are yet to be fully understood. However, it stands to reason that both allergen neutralization or Ab-mediated clearance could limit Ag availability for engagement with IgE on mast cells and basophils. In our studies, we found that serum titers of Ag-specific IgG2c were short-lived, however tertiary Ag exposure could re-boost serum IgG2c titers without any observed increase in IgE. Notably, the boosted IgG2c response occurred in the absence of continued IL-4Rα blockade illustrating a sustained reprogramming of the Ag-specific B cell response.

The molecular cues which sustain the MBC2 phenotype have not yet been described. In the T cell compartment, a subset of T helper cells, termed “Th2A” is uniquely associated with allergic disease, much like allergen-specific MBC2s (*68*). Th2A cells exhibit heightened Th2-like properties and are terminally differentiated, persisting even with IL-4Rα blockade (*18*, *68*, *69*). Here, we demonstrate that unlike Th2A cells, MBC2s do not harbor a refractory pathogenic phenotype. This requirement of IL-4 to sustain heightened CD23 and IL-4Rα expression was consistent amongst mouse models and humans and reaffirms early observations of IL-4-regulated expression in bulk B cell populations (*36*, *70*). Interestingly, CD23 purportedly augments allergen-specific responses through IgE-facilitated Ag-presentation (*71*, *72*). Thus, it is plausible that depressed CD23 expression observed not only on MBC2s, but throughout the B cell compartment, may curtail reactivation of allergen-specific CD4 T cells.

Our analysis of BCR sequences revealed clonal relationships between IgG2b/c and all upstream isotypes, including IgM/D, IgG3, and IgG1. These data, therefore, suggest that both naïve and Ag-experienced B cells can contribute to the A4RA-induced IgG2b/c response. Indeed, using molecular fate mapping we showed that when boosting with a type 1 adjuvant, Ag-specific IgG2c^+^ B cells can arise from GC-dervied IgG1^+^ MBCs that were produced during the primary type 2 response. Sequential CSR has been most extensively studied in the IgM to IgE trajectory, where IgG1 is a dominant intermediary. Interestingly, over 50% of the bulk IgG2 repertoire in humans also arises through sequential CSR with an IgG1 intermediary (*73*). This is somewhat surprising given that sequential rounds of V(D)J recombination are likely more metabolically taxing to the host. One explanation for this could be that sequential, rather than direct, CSR provides a means to regulate pro-inflammatory Ab-mediated effector functions. In agreement, both vaccination and allergy yield IgG1 dominant responses, though with chronic Ag exposure through repeated mRNA vaccination or allergen immunotherapy, a shift to the “anti-inflammatory” isotype, IgG4, is observed (*74–78*). Here, we demonstrated that Ag exposure in the absence in IL-4/IL-13 signaling could produce a similar effect – a deviation from the IgG1-IgE axis to an IgG2b/c response. As IgG2b/c is uninvolved in allergic inflammation, this shift to allergen-specific IgG2b/c production could be considered “anti-inflammatory”. Nevertheless, clonal relationships between all IgG subclasses and downstream IgE have been reported (*9*); whether the induced non-IgG1-expressing MBCs can later revert to a pathogenic IgE fate in humans is yet to be studied.

In summary, we propose the following model through which the IgE-fated B cell response is reshaped in the context of A4RA: 1) Allergic sensitization yields IgG1^+^ MBCs and type 2-polarized T cells, marked by upregulated IL-4 production. In an unperturbed recall response, IL-4 engages with IL-4R on IgG1^+^ MBCs enabling IgE CSR. 2) Administration of an IL-4R𝘢 blocking Ab prohibits IL-4 signaling which permits the emergence of a type 1 skewed cytokine response, marked by increased IFN-γ. 3) In this context, naïve and MBCs engage with cognate Ag in a type 1-skewed environment, favoring non-IgE CSR, as well as a loss of the type 2 MBC signature.

A limitation of our study is the disparities between mouse and human Ig isotypes, particularly the lack of clear IgG2b/c homologs. In humans, allergen desensitization is most often associated with increases in allergen-specific IgG2 and IgG4 (*75*, *79–81*). Both isotypes only weakly bind Fc receptors relative to other IgG subclasses and thus, may primarily regulate the IgE-mediated pro-inflammatory functions through competition for Ag binding (*82*, *83*). IgG2 is postulated to also regulate IgE effector functions directly through engagement with an inhibitory receptor, CD32b, on mast cells and basophils (*84*). Moreover, the cytokines that direct IgG subclass CSR differ between mice and humans. In mice, IFN-γ alone is necessary for IgG2c CSR, whereas in humans, it is suggested to work in concert with IL-6 and/or IL-2 to direct IgG CSR (*85*, *86*). Although human B cells do express the IFN-γ receptor, its direct signaling appears to mediate spontaneous GC formation and autoantibody production more so than CSR (*87*, *88*). Importantly, however, the MBC2 phenotype is present in both mouse and human allergen-specific IgG1^+^ B cells (*14*). Accordingly, even though the precise isotype output of a reprogrammed response otherwise fated to become IgE remains elusive, the diminished MBC2 phenotype provides promise of non-IgE CSR. This limitation underscores the importance of conducting parallel studies in humans to determine the reprogrammed fate of allergen-specific MBCs.

IL-4Rα blockade (dupilumab) is licensed for the treatment of various atopic diseases, such as moderate-to-serve asthma and chronic rhinosinusitis. Dupilumab significantly reduces serum IgE titers in these contexts, along with a decrease in allergen-specific IgE in trials involving food-allergic patients (*89–94*). Whether this treatment reprograms the pathogenic IgE-fated response has not yet been well addressed. A recent study uncovered that MBC2 frequency declines in atopic dermatitis patients treated with dupilumab, supporting the proposition that IL-4Rα blockade may alter the trajectory of IgE-fated MBCs (*38*). The product of an altered MBC2 trajectory in humans and if it is sustained following treatment cessation remains unknown.

In summary, our research demonstrates that IL-4/IL-13 signaling is required to sustain the IgE fate of B cell responses in established allergic disease and that loss of IL-4/IL-13 signaling can transform the allergen response to an innocuous type 1 phenotype. This not only bolsters the applicability of anti-IL-4Rα as adjunct therapy to other immunomodulatory interventions, but also broadly illuminates the potential to reprogram deviant B cell responses. Thus, further research is warranted to establish the intrinsic determinants that govern MBC fate, which will be imperative in the development of therapeutic strategies to correct deviant B cell responses in settings like allergy, autoimmunity, and transplantation.

## MATERIALS AND METHODS

### Mice

Female C57BL/6 mice (6-8 weeks old) were purchased from Charles River Laboratories (Wilmington, MA). Female B6.129S2-*Ighm^tmCgn^* and B6.SJL-*Ptprc^a^ Pepc^b^*/Boy mice (6-8 weeks old) were purchased from Jackson Laboratories (Bar Harbor, ME). *Igk^Flag/Strep^* and *S1pr2-cre^ERT2^* mice were a generous gift from Dr. Gabriel Victora (The Rockefeller University). All experimentation was in compliance with the McMaster University Research Ethics Board.

### Immunizations

*Epicutaneous sensitization:* Epicutaneous sensitization to ovalbumin (Grade V; Sigma-Aldrich, St. Louis, MO) was conducted as previously described (*21*). For recall responses, three low-back subcutaneous injections of 100 μg OVA were given over the course of a week. *Gastric sensitization:* Mice received one intragastric gavage of 1 mg OVA with 5 μg cholera toxin (CT) in PBS. Four weeks later, mice received 1 mg OVA in PBS weekly for three weeks via gavage. *Subcutaneous sensitization:* Mice were immunized subcutaneously with 250 μg OVA and 5 μg CT. *Intraperitoneal sensitization:* Mice were immunized intraperitoneally with 250 μg OVA and 250 μL Alhydrogel 2% (aluminum hydroxide [alum], InvivoGen, San Diego, CA), 40 μg CpG, or 5 μg CT.

Serum for analysis of OVA-specific antibodies was collected through retro-orbital bleeding. For allergen challenges, 5 mg OVA was intraperitoneally injected. Core body temperature and clinical signs were recorded, as previously described, and hematocrit was measured at 40 minutes (*95*).

### Antibody and Drug Administration

Anti-CD40L (MR-1), anti-IFN-γR1 (GR-20), and anti-trinitrophenol IgG2a isotype control (2A3) were purchased from Bio X Cell (Lebanon, NH). Anti-IL-4Rα (M1) was produced in-house, as previously described (*18*). Original hybridomas were generated by Immunex Corporation (Amgen, Thousand Oaks, CA) and were generously provided by Dr. Fred Finkelman (University of Cincinnati).

One mg of anti-IL-4Rα, anti-IFN-γR1, or isotype control was intraperitoneally injected the day prior to s.c. re-exposures. Anti-CD40L (0.35 mg) or isotype control was intraperitoneally injected on days -1, 0, and 2 of s.c. re-exposures.

200 µl of 50 mg/ml of tamoixfen (Sigma-Aldrich, Saint Louis, Missouri) was administrated by intragastric gavage on days 4, 8, and 12 post-immunization.

### Tetramer Staining and Enrichment

Purified OVA (Sigma-Aldrich, St. Louis, MO) was biotinylated (ThermoFisher Scientific, Waltham, MA) and tetramerized with streptavidin-PE or streptavidin-APC (Agilent, Santa Clara, CA), as previously described (*31*, *32*). For decoy tetramer, β-lactoglobulin (Sigma-Aldrich, St. Louis, MO) was biotinylated. Streptavidin-PE or -APC was conjugated with DyLight 594 or DyLight 755 esters (ThermoFisher Scientific) and was used to tetramerize the biotinylated-β-lactoglobulin.

Cell pellets were resuspended and stained with 5 nM decoy tetramer in 20 μL FACS buffer (10% FBS and 1 μM EDTA in PBS) and 2% rat serum for five minutes on ice. Five nM OVA tetramer was added and incubation continued for 30 minutes. Cells were washed to remove unbound tetramer and were incubated with anti-PE or -APC microbeads (Miltenyi Biotec, Germany) for 15 minutes on ice. Cells were passed through LS columns affixed to MACS magnetic separator (Miltenyi Biotec). Bound fractions were eluted with FACS buffer.

For CD45.1 enrichments, cell pellets were resuspended in 20 μL FACS buffer with 0.25 μg FcBlock (BioLegend) for five minutes on ice, then stained with anti-CD45.1-APC (A20; BioLegend) for 30 minutes. Enrichment with anti-APC microbeads proceeded as described above.

### Patient Samples

A cohort of three peanut allergic donors, aged 9-23, were recruited to donate blood from McMaster Children’s Hospital and Hamilton Health Sciences (Hamilton, Ontario, Canada). Allergic status was confirmed by peanut-specific IgE (higher than 0.35 kU/L) ImmunoCap at Laboratory Reference Centre Hamilton (McMaster Children’s Hospital). Donors were excluded from the study for previous or current allergen immunotherapy, omalizumab, dupilumab or other systemic immunomodulatory treatment, as well as diagnosed autoimmune or immunodeficiency diseases. All donors were recruited with written consent and ethical approval from the Hamilton Integrated Research Ethics Board.

Peripheral blood was collected into heparinized tubes (BD). PBMCs were isolated via Ficoll-Paque (GE Healthcare) density centrifugation and were cryopreserved in FBS + 10% DMSO for long-term storage in liquid nitrogen. Before culture, viability of thawed PBMCs was confirmed to be ≥ 70%. PBMCs were cultured for 8 days and stimulated with 2.5 ng/mL crude peanut extract (Stallergenes Greer) with or without 50 ug/mL anti-human IL-4Rα (MAB230; R&D Systems), as previously described (*18*).

### Flow Cytometry and Cell Sorting

Single-cell suspensions were plated in U-bottom 96-well plates and incubated with 0.25 μg FcBlock for 10 minutes on ice. Cells were stained extracellularly for 30 minutes on ice with Ab cocktails prepared in FACS buffer. For intracellular staining, cells were fixed and permeabilized with Cytofix/Cytoperm (BD Biosciences), as per manufacturer’s instructions. Intracellular Ab cocktails were prepared in 1x perm wash and cells were stained for 30 minutes on ice.

The following anti-mouse fluorochrome-conjugated antibodies were used: B220-AF700 (RA3-6B2), CD138-PE/Dazzle 594 (281–2), CD3-BV711 (17A2), CD3-BV510 (17A2), CD45.1-APC (A20), F4/80-BV711 (BM8), F4/80-BV510 (BM8), CD38-PE/Cy7 (90), GL7-PerCP.Cy5.5 (GL7), GL7-BV421 (GL7; BD, Franklin Lakes, NJ), IgD-Pacific Blue (11-26c.2a), IgD-BV605 (11-26c.2a), IgM-BV786 (II/41; BD), IgM-FITC (II/41), IgG1-BV650 (RMG1-1), IgG2b-PECy7 (RMG2b-1), IgG2c-FITC (polyclonal; SouthernBiotech, Birmingham, AL), IL-4Rα-PE (REA235; Miltenyi Biotec), CD23-BV421 (B3B4), Flag-APC (L5), Anti-Streptavidin-Biotin (5A9F9; LSBio, Newark, CA), and Streptavidin-BV421 (405225). The following anti-human antibodies were used: IL-4Rα-PE (G077F6), CD23-BV421 (EBVCS-5), IgD-AF700 (IA6-2), CD27-BV605 (O323), IgM-BV650 (MHM-88), CD38-PECy7 (HB-7), CD19-PerCP.Cy-5.5 (SJ25C1), CD3-BV711 (OKT3), CD14-BV711 (M5E2), and CD16-BV711 (3G8). All antibodies were purchased from BioLegend (San Diego, CA), unless otherwise noted. Viability staining was executed with Fixable Viability Dye eFluor780 (eBioscience, San Diego, CA) or Fixable Viability Stain 510 (BD Biosciences).

Data were acquired with an LSRFortessa flow cytometer (BD). Bulk cell sorting was performed on a MoFlo XDP cell sorter (Beckman Coulter, Brea, CA). All data were analyzed with FlowJo, version 10 software (Tree Star, Ashland, OR).

### Adoptive Transfers

Spleens and mesenteric LNs were collected four weeks following intraperitoneal sensitization of CD45.1^+^ mice with OVA and Alum. Class-switched MBCs (B220^+^ CD3^-^ F4/80^-^ CD38^+^ GL7^-^ IgD^-^ IgM^-^) from tetramer enriched fractions were isolated by flow sorting. Ten thousand cells were intravenously injected in the tail-vein of naïve recipient (muMT) mice. The day after transfer recipients were intraperitoneally injected with 250 μg OVA with either 40 μg CpG or 5 μg CT. One week later, spleens and mesenteric LNs were collected for flow cytometric analysis of donor B cells, and serum was collected for OVA-specific Ig analysis by ELISA.

### ELISAs

OVA-specific Igs were measured by sandwich ELISA, described previously, with some modifications (*96*). Briefly, MaxiSorp plates (ThermoFisher Scientifc) were coated with 100 ng anti-IgG2b, -IgG2c, -IgE, or -IgA overnight at 4°C. Plates were blocked in 5% skim milk at 37°C for one hour, then diluted (1:2) samples were added and incubated overnight at 4°C. Digoxigenated OVA (15 ng) prepared in-house was added for two hours at room temperature, followed by anti-digoxigenin Fab fragments (Roche, Basel, Switzerland) for one hour. Plates were developed with 3,3’,5,5’-tetramethylbenzidinie (Sigma-Aldrich) and stopped with sulfuric acid. Optical densities were acquired with a Multiskan FC photometer (ThermoFisher Scientific).

For OVA-IgG1, plates were coated with 400 ng OVA in carbonate-bicarbonate buffer overnight at 4°C. Plates were blocked in 5% skim milk at 37°C for two hours, then diluted (1:20) samples were added and incubated overnight at 4°C. Biotinylated anti-mouse IgG1 (SouthernBiotech; 1.25 ng) was added for two hours at room temperature, followed by alkaline-phosphatase streptavidin (ThermoFisher Scientific) for one hour. Plates were developed with 4-nitrophenyl phosphate (Sigma-Aldrich) and stopped with sodium hydroxide. Optical densities were acquired as above.

### 10x Chromium Sequencing & Analysis

#### Sequencing

Spleens and inguinal LNs were collected one month post-recall response from epicutaneously or subcutaneously sensitized mice and B cells were sorted for sequencing. Single-cell 5’ gene expression and V(D)J libraries were prepared with Chromium Single-Cell V3 Reagent Kit and Controller (10x Genomics; Pleasanton, CA). Libraries were sequenced with NextSeq2000 (Illumina; San Diego, CA). Data were processed with the Cell Ranger Count and V(D)J pipelines (version 7.0.1).

#### Gene Expression Analysis

Sequencing reads were aligned to the mouse transcriptome (GRCm38), followed by filtering, and correction of cell barcodes and Unique Molecular Identifiers (UMIs). Reads associated with retained barcodes were quantified and used to build expression matrices for further analysis with Seurat (version 4.1.3). Cells expressing less than 200 genes, genes that were detected in less than 3 cells, and cells expressing over 10% mitochondrial genes were filtered out. Resulting cell counts per condition were CT = 3016, IC = 2478, and A4RA = 1875. Data from separate experimental conditions were integrated using reciprocal PCA. Normalization, variable feature selection, and scaling were performed with Seurat default parameters. Principal component analysis (PCA) was performed and the top 15 components were used for UMAP dimensionality reduction and clustering at resolution 0.4. Contaminating non-B cells were filtered out and PCA and clustering were repeated on the subsetted Seurat object at resolution 0.3. Differentially expressed genes or gene modules (GO:0006119, R-MMU-983705, GO:0034340, GO:0071559, GO:0034341, GO:0070670) were identified with the Wilcoxon Rank Sum test.

#### BCR Analysis

BCR analysis was performed with pipelines in the Immcantation suite (*64*). The “filtered_contig.fasta” and “filtered_contig_annotations.csv” outputs from Cell Ranger V(D)J were used to assign V, D, and J genes with IgBLAST and cells assigned with 0 or >1 heavy chains were removed. Clonal groups were defined with the hierarchialClones function using applying a Hamming distance of 0.1. Germline sequences were reconstructed with createGermlines and IMGT germline references (version 3.1.39). SHM frequency within the IGHV gene was calculated with observedMutations. BAM files from the Cell Ranger output were used to map sterile *Igh* transcripts with the sciCSR pipeline (*97*).

### Bulk VDJ Sequencing & Analysis

#### Sequencing

Spleens and inguinal LNs were collected one month post-recall response from epicutaneously sensitized mice and OVA^+^ B cells were sorted directly into Buffer RLT (Qiagen; Hilden, DE) prior to shipment on dry ice to iRepertoire. RNA extraction was performed prior to 3-chain long read library preparation (Mouse BCR Heavy, Kappa, Lambda). Amplified PCR products were run on MiSeq System with MiSeq Reagent Nano Kit v2 (iRepertoire, Huntsville, AL).

#### Analysis

Nucleotide sequences and UMI count were extracted from “joinedSeq” and “copy” columns of the “pep.csv” output from iRweb (iRepertoire) and were aligned with IgBLAST (version 1.18.0) using IMGT reference sequences. Processing of IgBLAST output was performed with the Immcantation suite to assign V, D, and J genes, cluster into clonal groups, assign unmutated germlines, and calculate IGHV SHM hypermutation frequency, as described in the 10x Chromium section. Lineage tree topologies were predicted by maximum parsimony in IgPhyML with constraints for unidirectional isotype switching and trees were subsequently visualized with dowser (*98*, *99*).

### Smart-Seq2 Sequencing & Analysis

#### Sequencing

Spleens and inguinal LNs were collected one month post-recall response from epicutaneously sensitized mice. OVA-specific B cells were-single cell sorted into 96 U-bottom well PCR-grade RNase and DNase free plates before being flash frozen at −80C. Full-length cDNA was generated with Smart-Seq Single Cell Kit (Takarabio; Mountain View, CA). Libraries were then prepared for sequencing with NovaSeq 600 Reagent Kits (Illumina; 20028400) and sequenced on Illumina NovaSeq 6000 (Ilumina; San Diego, CA).

#### Analysis

Sequencing reads were aligned to the mouse transcriptome (GRCm38) with STARsolo (*100*). Count matrix output was normalized and scaled with Seurat (version 4.1.3). Datasets from separate sequencing runs were integrated using reciprocal PCA. BCR sequences were reconstructed with the TRUST4 algorithm to obtain the productive isotype for each cell (*101*, *102*). BAM files from the STARsolo output were then used to map sterile *Igh* transcripts with the sciCSR pipeline (*97*).

### Statistical Analysis

Statistical analyses not pertaining to RNA sequencing were performed with Prism 9 (GraphPad Software; San Diego, CA). Comparisons were drawn between groups (CT, IC, A4RA) or across time points using two-sided Student’s *t*-test or ANOVA. Data were considered significant with a *p* value < 0.05.

## Acknowledgments

We kindly thank S. Eisenbarth for thoughtful review of the manuscript. We also thank M. Subapanditha for assistance with cell sorting and C. Mader, the McMaster Genomics Facility, and G. Kongsgaard Koed, and J. Skovsgaard for sequencing support. We thank J. Ng for guidance with sciCSR pipeline. Experimental schematics were created with BioRender (www.biorender.com). KB was supported by a CIHR CGS-Doctoral award and CIHR Postdoctoral Fellowship. AP was supported by the Schroder Allergy & Immunology Research Institute Scholarship and an Ontario Graduate Scholarship.

## Funding

Schroeder Foundation (SW, MJ)

Food Allergy Canada (SW, MJ)

ALK Abelló (SW, MJ)

Canadian Allergy, Asthma, and Immunology Foundation (MJ)

Michael Zych Family (SW, MJ)

## Author contributions

Conceptualization: KB, AP, MJ

Investigation: KB, AP, AA, AF, TDW, JW, SSK-B, DDL, OM-D, SG, EG, JFEK, AG

Visualization: KB, AP

Funding acquisition: SW, PSA, MJ

Supervision: MJ

Writing – original draft: KB, AP

Writing – review & editing: KB, AP, MJ

## Data and materials availability

This manuscript produced no new code. Single-cell and bulk sequencing data will be deposited to the GEO database prior to publication. All other data are available within the main text or the supplementary materials.

---

**Fig. S1.**
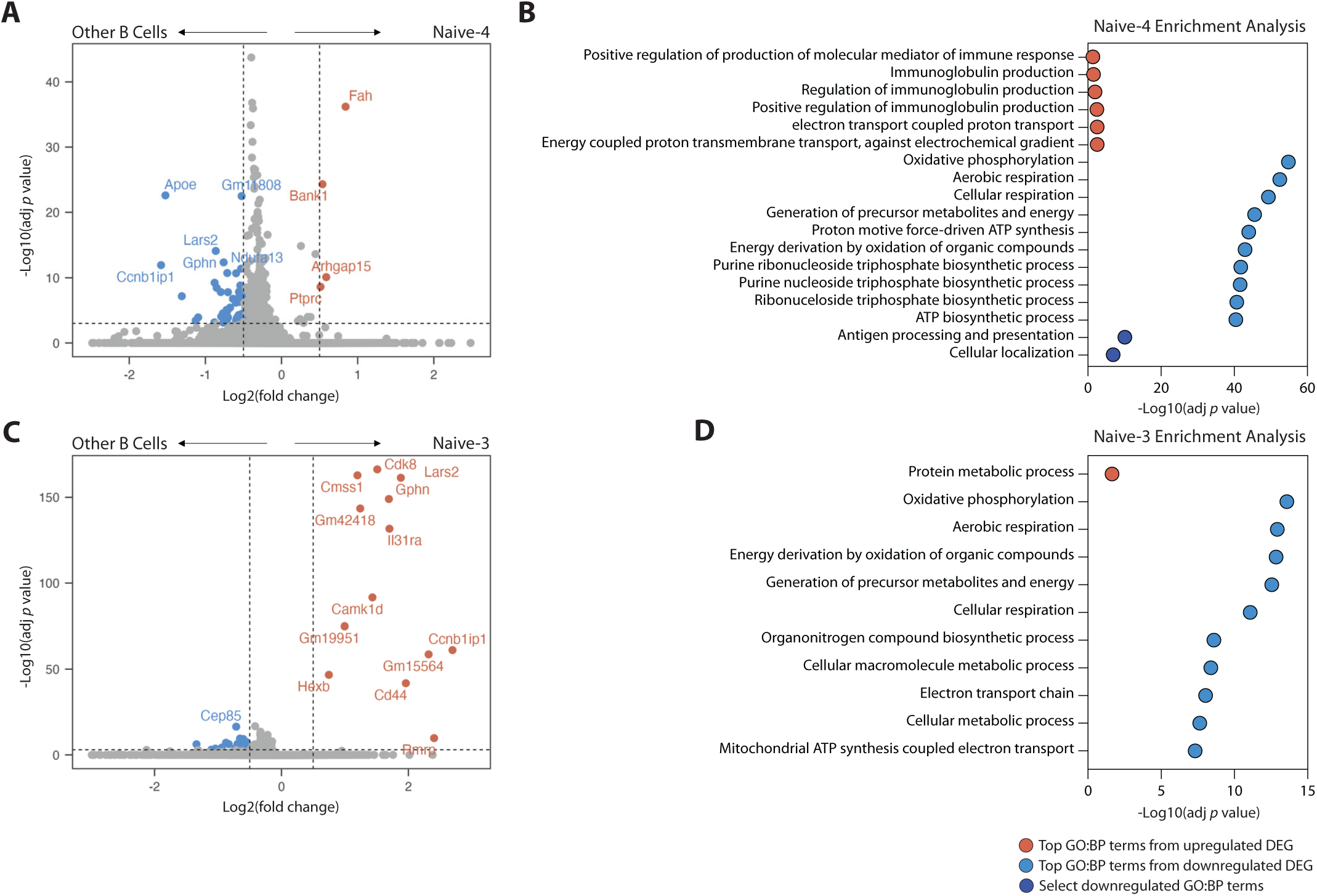
Gene expression profiles of naïve B cell clusters. **(A)** Volcano plot of differentially expressed genes in Naïve-4 (A) and Naïve-3 (C) clusters. Red points indicate genes with -log10 adjusted *p* value > 5 and log2 fold change ± 0.5. **(B, D)** GO:BP enrichment analysis of significantly upregulated (red) and downregulated (blue) genes, excluding ribosomal genes, in Naïve-4 (B) and Naïve-3 (D) clusters. DEG, differentially expressed genes.

**Fig. S2.**
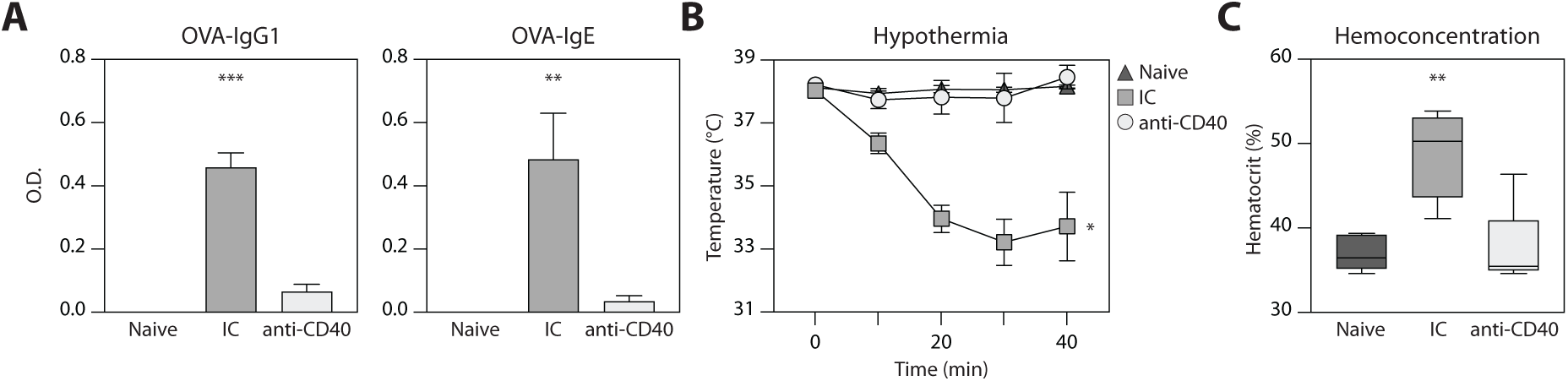
Requirement of B-T interactions for IgE recall responses. **(A)** Serum OVA-specific immunoglobulins immediately prior to allergen challenge. **(B)** Core body temperature. **(C)** Hemoconcentration measured 40-minutes post-allergen challenge. Data represent 2 experiments, each with 4 mice per group. Data are presented as mean ± SEM (A, B) or median ± min/max (C). * *p* < 0.05, ** *p* < 0.01, *** *p* < 0.001.

**Fig. S3.**
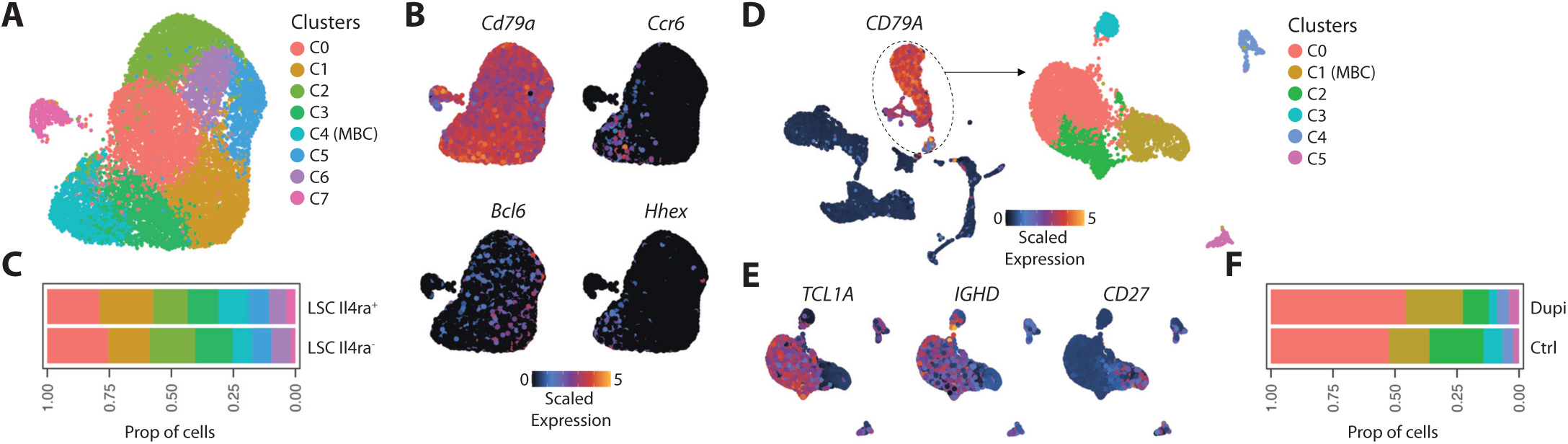
Analysis of publicly reposited scRNA-seq datasets. **(A, D)** UMAP of B cell clusters from scRNA-seq analysis of Duan & Liu *et al.* (2021) (A) and Mountz & Gao *et al.* (2022) (D) datasets. **(B, E)** UMAP plots of scaled gene expression used to define MBC clusters. **(C, F)** Percent of cells in each cluster. LSC, lymphoid stromal cell.

**Fig. S4.**
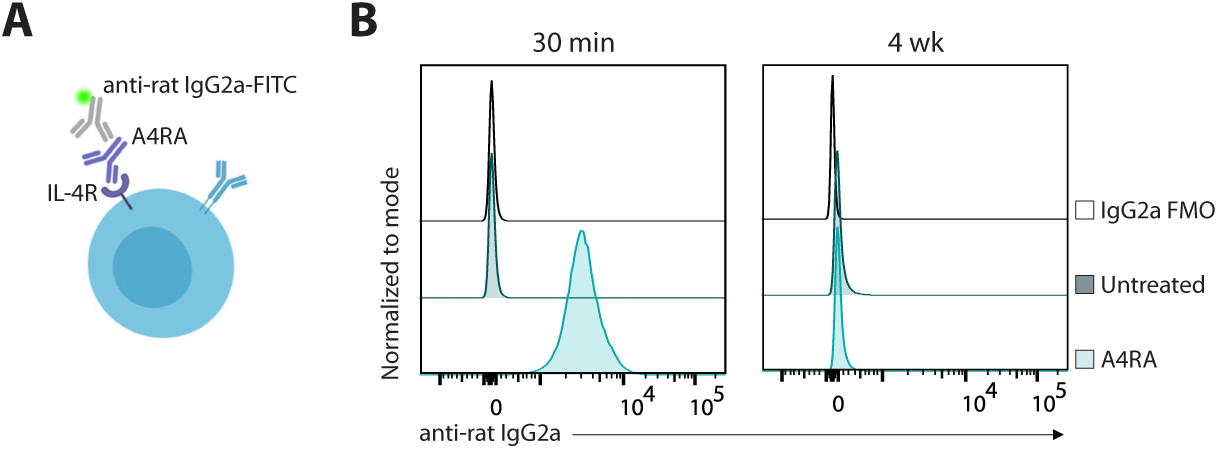
Surface expression of A4RA blocking antibody. **(A)** A4RA-treated cells were stained with an anti-rat IgG2a-FITC antibody to detect the Fc region of A4RA. **(B)** Representative histograms of surface rat-IgG2a expression in mouse B cells four weeks after *in vivo* A4RA administration (right). As a comparison, splenocytes from untreated mice were incubated with A4RA for 30 minutes prior to staining with fluorochrome conjugated antibodies (left). FMO, fluorescence minus one.

**Fig. S5.**
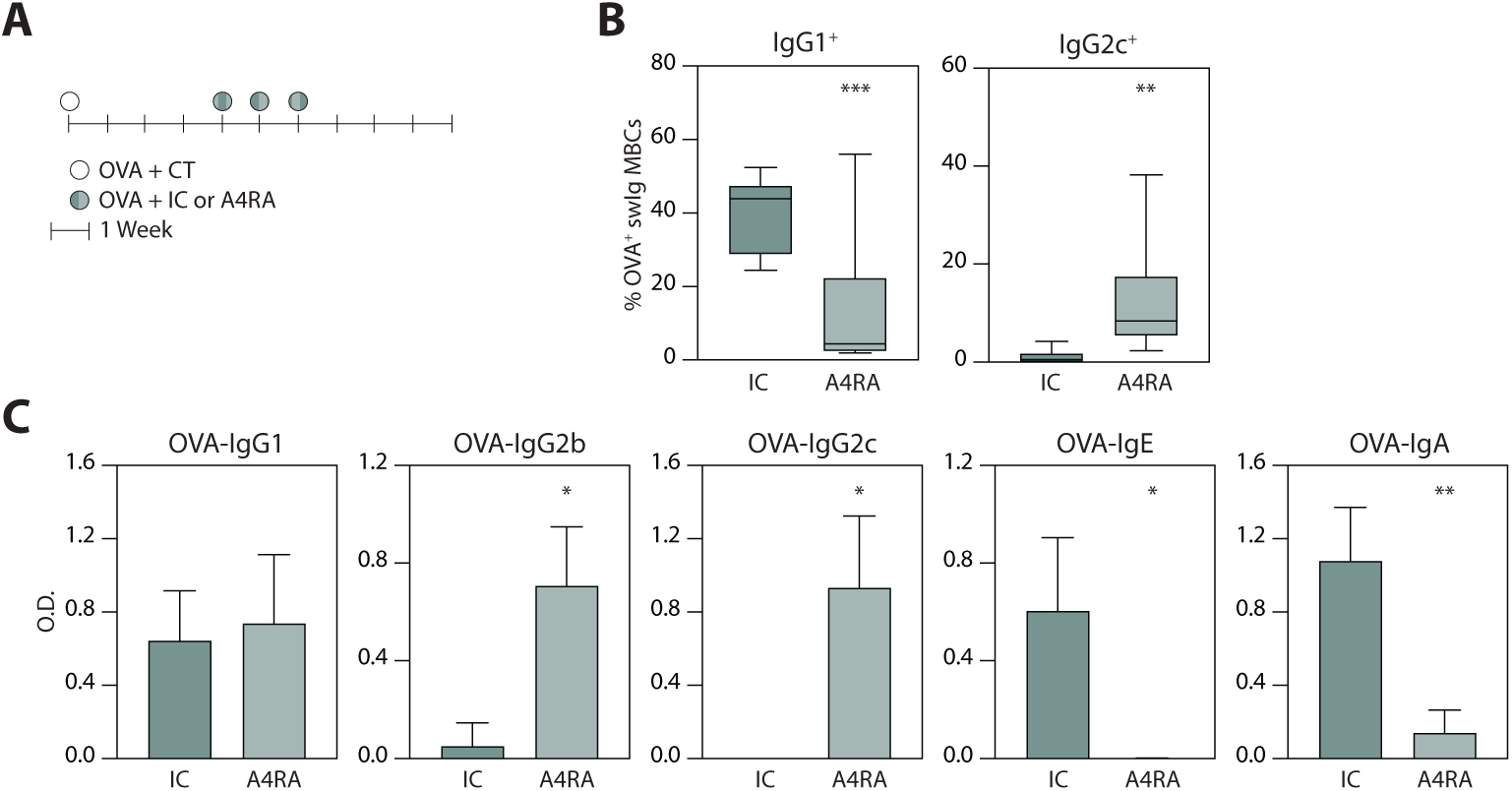
B cell reprogramming in orally-sensitized mice. **(A)** Experimental schematic. **(B)** Summary plots of IgG1+ (left) and IgG2c+ (right) MBC frequency oral re-exposures. **(C)** OVA-specific serum immunoglobulin profiles post-boost. Data represent 2 experiments, each with 4 mice per group. Data are presented as median ± min/max (B) or mean ± SEM (C). * *p* < 0.05, ** *p* < 0.01, *** *p* < 0.001.

**Fig. S6.**
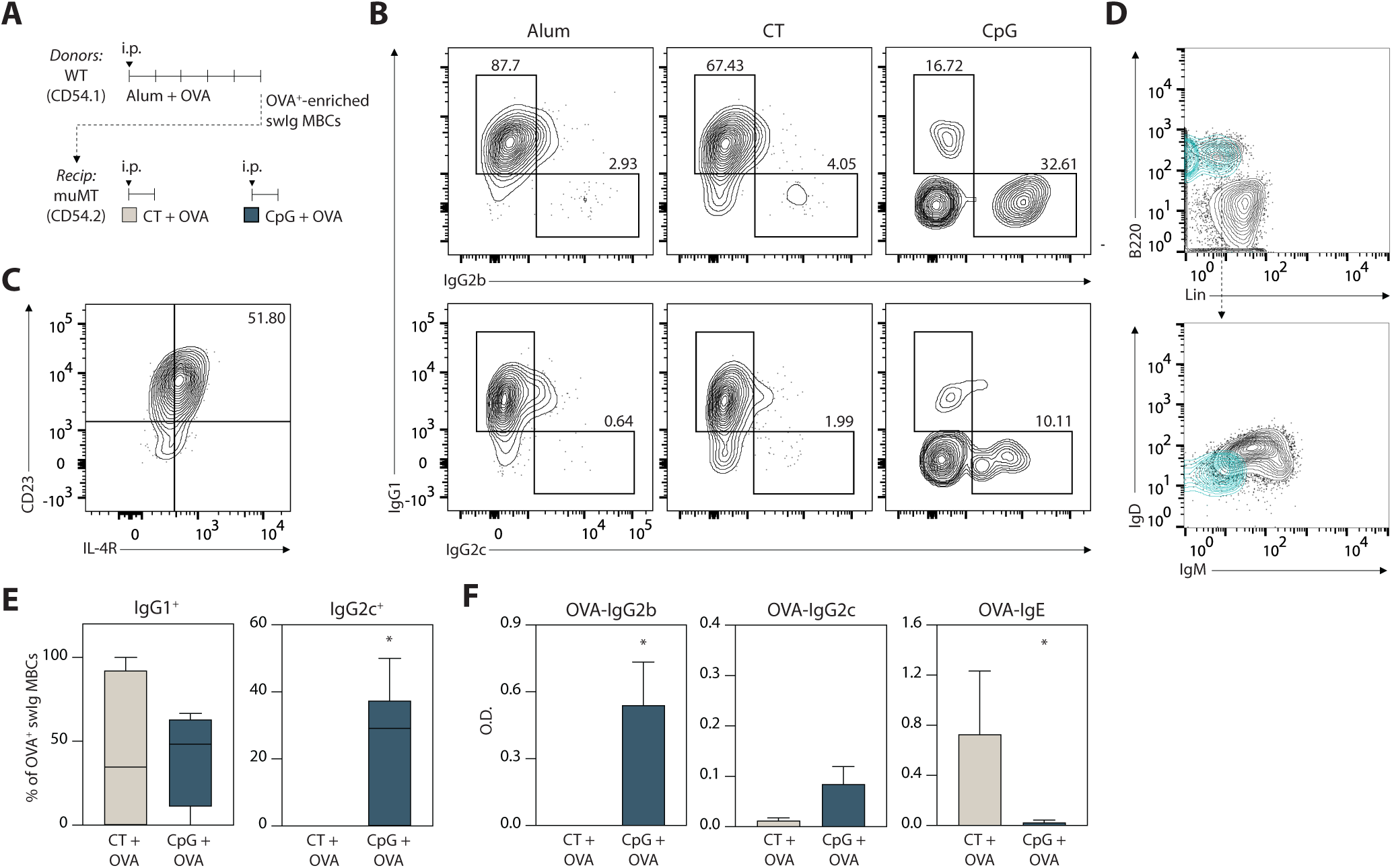
Boost with CpG produces IgG2 response from type 2-primed class-switched MBCs in adoptive transfer model. **(A)** Adoptive transfer schematic. **(B)** To illustrate the isotype distribution of class-switched MBCs, WT mice were immunized with OVA + Alum, CT, or CpG. Representative contour plots class-switched OVA+ MBCs. Pre-gated as OVA^+^ B cells > CD38^+^ GL7^-^ > IgM^-^ IgD^-^. Numbers indicate the proportion of IgG1^+^, IgG2b^+^, and IgG2c^+^ cells from the parent class-switched MBC population. **(C)** Representative contour plot of CD23 and IL-4R expression from OVA^+^ IgG1^+^ MBCs in OVA + Alum-immunized mice. Number indicates the proportion of CD23^+^ IL-4R^hi^ cells from the parent OVA^+^ IgG1^+^ MBC population. **(D)** Contour plots depicting the adoptively transferred class-switched B cell population (blue) from OVA-enriched B cells of OVA + Alum-immunized mice. **(E)** Summary plots depicting donor (CD45.1+) OVA+ IgG1+ and OVA+ IgG2c+ B cell frequency in recipient mice one week post-transfer and boost. **(F)** OVA-specific serum immunoglobulin profiles in recipient mice one week post-transfer and boost. Data represent 2 experiments, each with 2-4 mice per group. Data are presented as median ± min/max (E) or mean ± SEM (D, F). * *p* < 0.05.

**Fig. S7.**
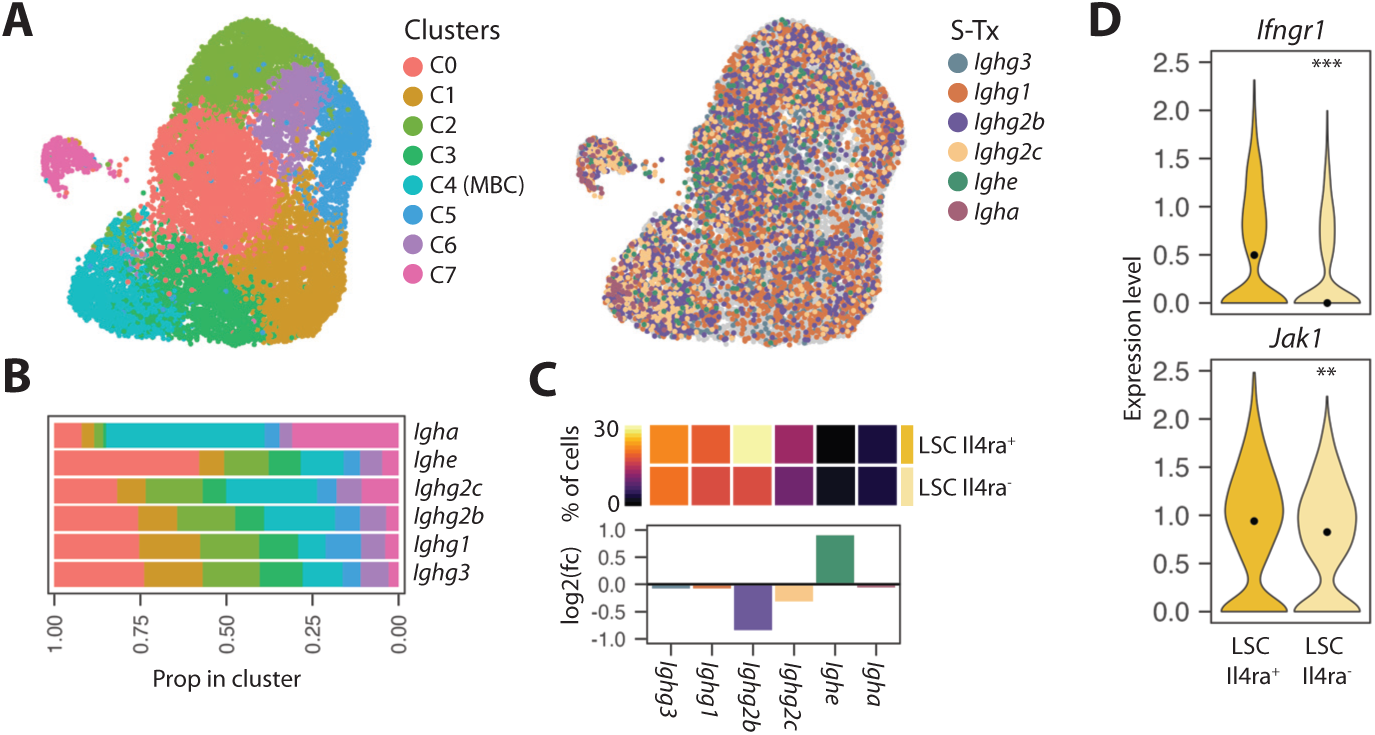
Ig S-Tx usage with increased IL-4 bioavailability in mice. **(A)** UMAP of B cell clusters from scRNA-seq analysis of Duan & Liu *et al.* (2021) and overlay of S-Tx expression (right). Colored points indicate the most downstream *Igh* S-Tx expressed by a cell. **(B)** Distribution of S-Tx amongst B cell clusters. **(C)** Heatmap displaying the percent of B cells expressing S-Tx downstream to their productive isotype and barplots depicting the fold change in S-Tx expression in LSC Il4ra^-^ compared to LSC Il4ra^+^. **(D)** Expression level of genes in MBC cluster. ** *p* < 0.01, *** *p* < 0.001.

**Fig. S8.**
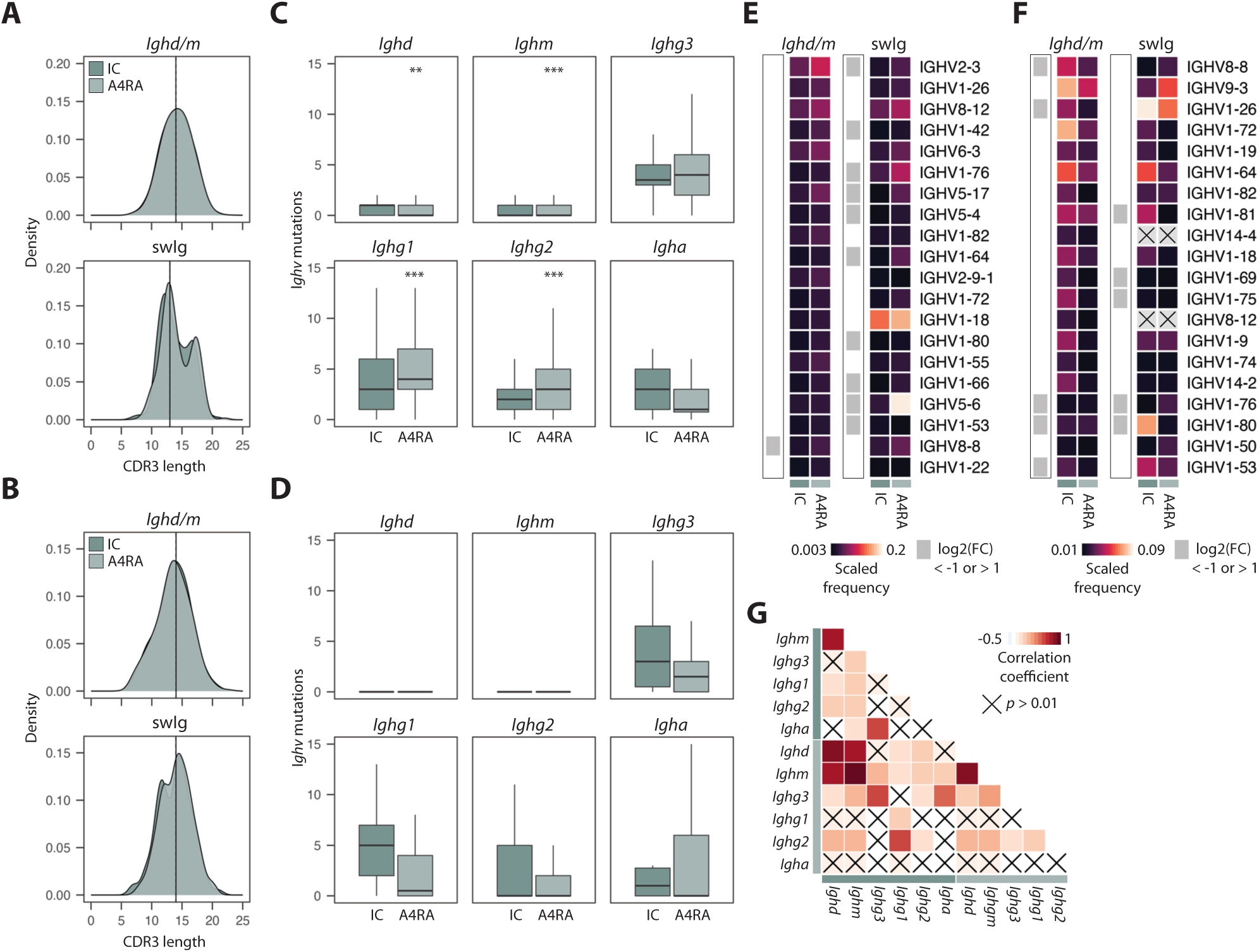
VDJ repertoire analysis in OVA-enriched bulk sequencing and 10X datasets. **(A, B)** Distribution of CDR3 amino acid length in OVA-specific B cells (A) and polyclonal B cells (B). Solid and dashed lines indicate median length for IC and A4RA, respectively. **(C, D)** SHM count in IGHV region of OVA-specific B cells (C) and polyclonal B cells (D) presented as median ± min/max. **(E, F)** Heatmap of scaled IGHV gene usage for top 20 used IGHV genes in OVA-specific B cells (E) and polyclonal B cells (F). Grey squares indicate IGHV genes with log2 fold change ± 1 between IC and A4RA. Crosses indicate IGHV genes that are not present in the swIg cluster. **(G)** Correlation matrix of paired IGHV-IGHJ gene usage between isotypes in IC and A4RA. ** *p* < 0.01, *** *p* < 0.001.

